# Expanding functional protein sequence space using generative adversarial networks

**DOI:** 10.1101/789719

**Authors:** Donatas Repecka, Vykintas Jauniskis, Laurynas Karpus, Elzbieta Rembeza, Jan Zrimec, Simona Poviloniene, Irmantas Rokaitis, Audrius Laurynenas, Wissam Abuajwa, Otto Savolainen, Rolandas Meskys, Martin K. M. Engqvist, Aleksej Zelezniak

## Abstract

*De novo* protein design for catalysis of any desired chemical reaction is a long standing goal in protein engineering, due to the broad spectrum of technological, scientific and medical applications. Currently, mapping protein sequence to protein function is, however, neither computationionally nor experimentally tangible ^1,2^. Here we developed ProteinGAN, a specialised variant of the generative adversarial network ^3^ that is able to ‘learn’ natural protein sequence diversity and enables the generation of functional protein sequences. ProteinGAN learns the evolutionary relationships of protein sequences directly from the complex multidimensional amino acid sequence space and creates new, highly diverse sequence variants with natural-like physical properties. Using malate dehydrogenase as a template enzyme, we show that 24% of the ProteinGAN-generated and experimentally tested sequences are soluble and display wild-type level catalytic activity in the tested conditions *in vitro*, even in highly mutated (>100 mutations) sequences. ProteinGAN therefore demonstrates the potential of artificial intelligence to rapidly generate highly diverse novel functional proteins within the allowed biological constraints of the sequence space.

## MANUSCRIPT

A protein’s three-dimensional structure, physicochemical properties and molecular function are defined by its amino acid sequence. From the 20 commonly occurring proteinogenic amino acids, a small sized protein comprising 100 amino acids can be made in 10^130^ unique ways. In this vast multidimensional space - often referred to as the protein fitness landscape ^4^ - as little as 1 in 10^77^ sequences are estimated to fold into the defined three-dimensional structures to carry out specific functions ^5–7^. This imposes a great burden on experimental approaches aiming to design novel protein sequences, such as random mutagenesis ^4^ and recombination of naturally occurring homologous proteins ^8,9^, as up to 70% of random amino acid substitutions typically result in a decline of protein activity and 50% are deleterious to protein function ^4,10–16^. On the other hand, Artificial intelligence (AI) is not limited by the amount of sequence variations it can process ^17–19^ and, instead of depending on a blind search, is based on an inference-based process - it infers protein properties ^18,20^ and function ^19,21^ directly from training examples. Recent AI approaches have also demonstrated great potential in capturing both the structural and evolutionary information found in natural protein sequences ^17,22^. Nevertheless, the majority of existing machine learning models in biology are *discriminative* ^17,18,21,23^, i.e., the model is trained using readily available data, to predict the properties of a given protein sequence. A *generative* modeling approach, in contrast, could generate new sequence samples from the learned portion of functional protein sequence space, providing direct access to unexplored sequence diversity within functional protein structures, without the need to test a large number of non-functional protein sequence variants. Indeed, breakthrough generative learning strategies, such as Generative Adversarial Networks (GANs) ^3^ can learn multidimensional distributions in disparate scientific domains to generate photorealistic images ^24^, hand-written text ^25^, music ^26^ and even DNA sequences with biological properties ^27,28^. Hence, in the present study we develop and test ProteinGAN (Figure 1a) to learn and sample the protein sequence space for *de novo* generation of functional enzymes.

**Figure 1.**
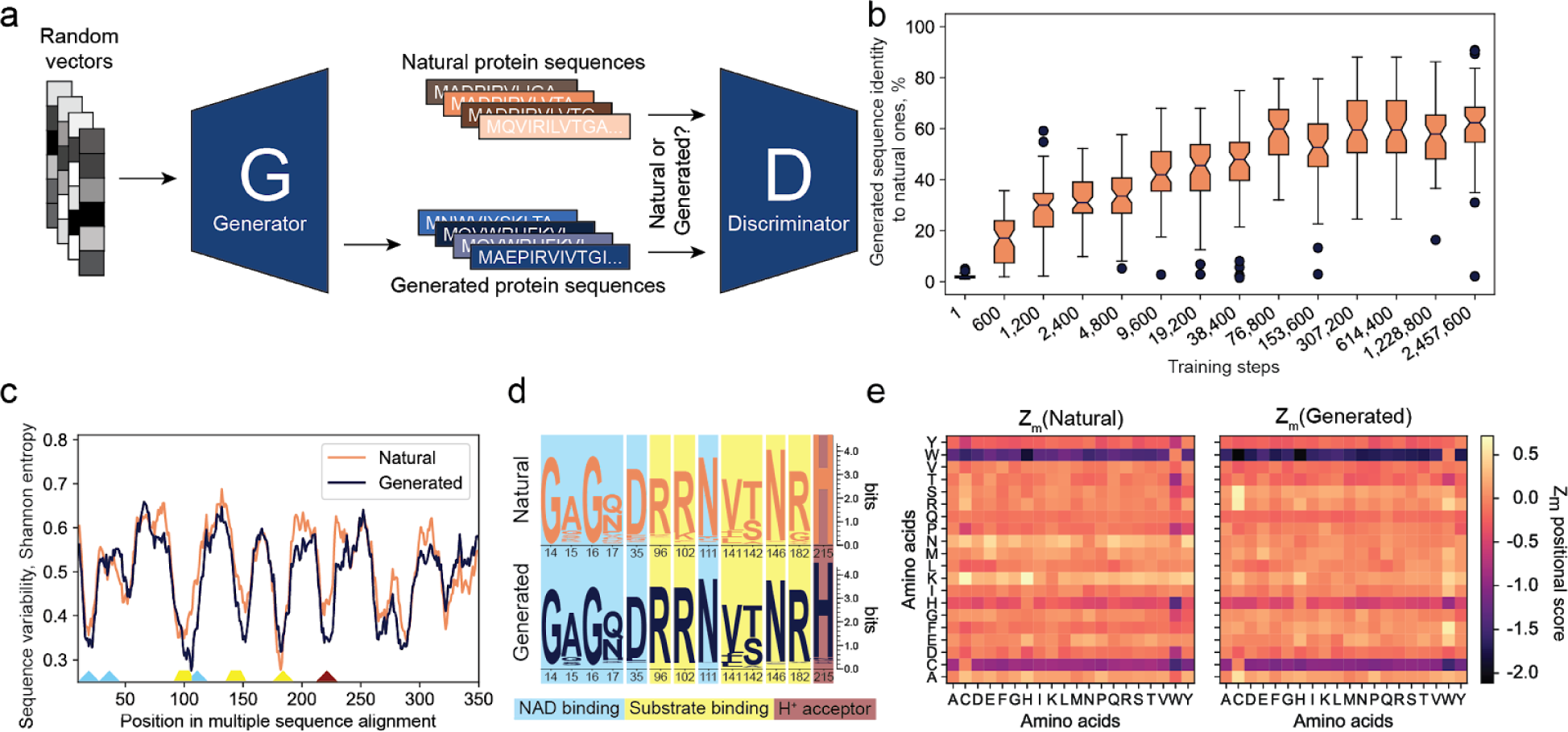
ProteinGAN learns intrinsic relationships of natural protein sequences. **a)** ProteinGAN training scheme. Given a random input vector, the Generator network produces a protein sequence which is scored by the Discriminator network comparing it to the natural protein sequences. The generator tries to fool the discriminator by generating sequences that will eventually look like real ones (the generator never actually sees real enzyme sequences). **b)** Sequence identity of 64 generated sequences to the nearest natural sequence at different training timestamps. **c)** ProteinGAN effectively captures amino acid distribution of natural MDH sequences. Sequence variability expressed as Shannon entropies for generated and training sequences estimated from multiple-sequence alignment (MSA). Low Shannon entropy values represent highly conserved and thus functionally relevant positions, whereas high entropy indicates high amino acid diversity at a given position. **d)** A sequence logo of key conserved positions in the multiple sequence alignment. **e)** ProteinGAN learns the order of amino acids in natural MDH sequences. Amino acid pair association (Z_m_ positional score) matrices for Natural and Generated protein sequences. Positive values indicate a larger distance than expected when comparing random sequences with the same amino acid frequency. The numbers indicate how many positions, on average, the amino acids in a pair are closer (negative values) or further apart (positive values) than in a random sequence.

ProteinGAN is a generative model architecturally tailored specifically to learn patterns in long biological sequences (Methods, Supplementary Figure 1). A customized temporal convolutional network ^29^ enabled the network to simultaneously analyze local and global sequence fragments, and thus to capture the meaningful sequence motifs and long-distance relationships that are critical for correct protein structural assemble. Additionally, we introduced a self-attention layer ^30^, to help ProteinGAN focus on functionally important areas across the entire length of the sequences, such as catalytic residues (Methods, Supplementary Figure 1). The final architecture of the network comprised 45 layers with over 60 million trainable parameters.

A family of bacterial malate dehydrogenase (MDH) enzymes (EC 1.1.1.37) was used to train the neural network. MDH is a tricarboxylic acid cycle enzyme catalyzing the conversion of malate to oxaloacetate using NAD^+^ as a cofactor (Supplementary Figure 2). We chose MDH based on the following criteria: (i) it has a large number of diverse sequences (a total of 16 706 unique sequences were used for training), which were on average 319 ± 18.2 (sd) amino acids long with pairwise sequence identities as low as 10%; (ii) it is a complex enzyme that must bind both its substrate and the NAD^+^ cofactor for catalysis and (iii) its activity can be readily monitored *in vitro*. During model training, we assessed the quality of the sequences generated by ProteinGAN at every 1200 learning steps. In each assessment, 64 sequences were generated and their identities to natural sequences in the training and validation datasets were computed (Figure 1b). After 2.5M learning steps, at which training was terminated (Supplementary Figures 3, 4), the mean sequence identities between the generated and natural sequence sets had reached a plateau (median sequence identity to the closest natural sequences was 61.3%). Following the initial quality assessment, 20 000 of the generated sequences were used to further evaluate the ProteinGAN performance.

**Figure 2.**
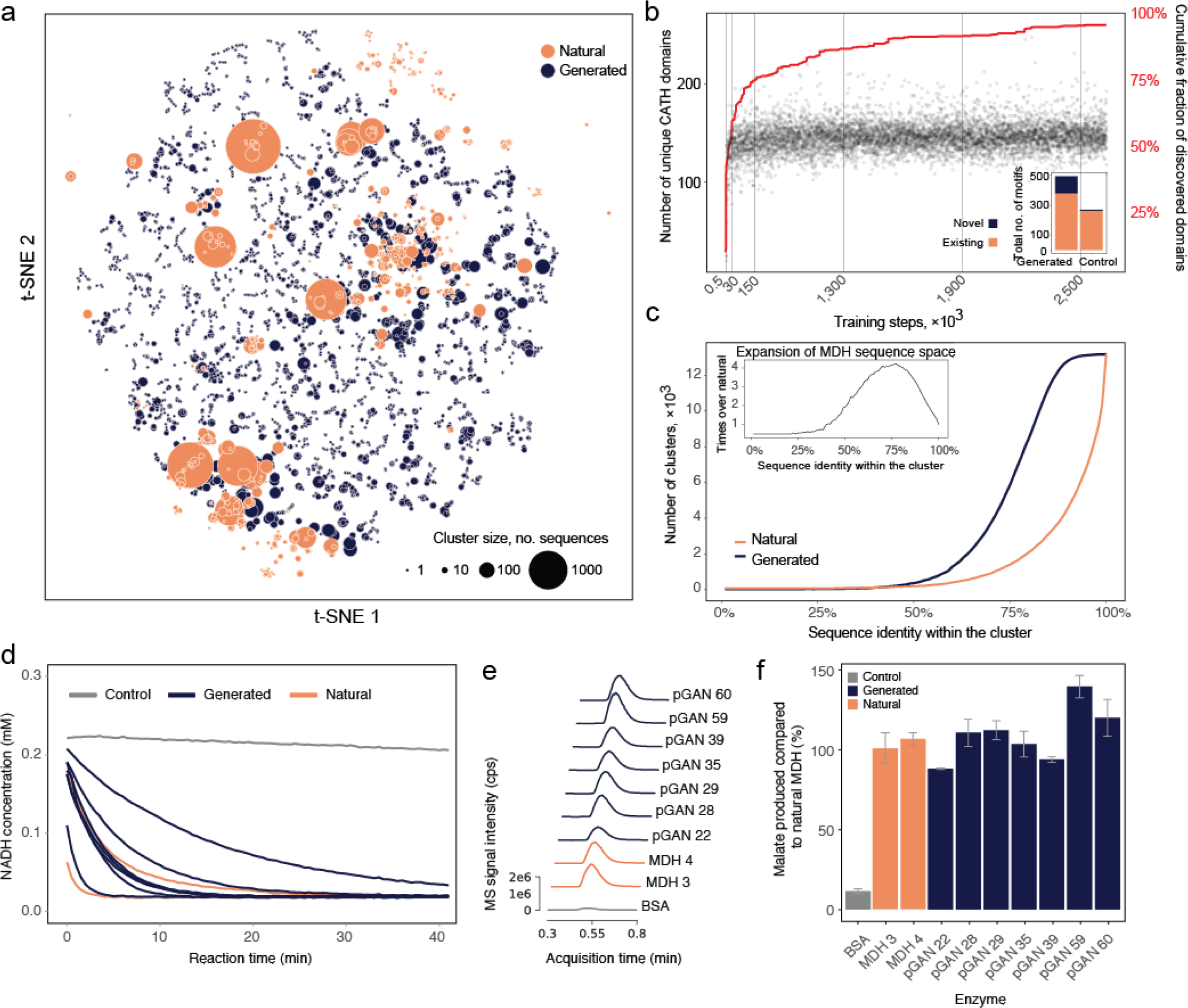
ProteinGAN expands the functional MDH sequence space. **a)** The protein sequence space was visualized by transforming a distance matrix derived from k-tuple measures of protein sequence alignment (Wilbur WJ, Lipman DJ. 1983) into a t-SNE embedding. Dot sizes represent the 70% identity cluster size for each representative. As opposed to natural sequences, generated sequences formed disparate small clusters indicating their diverse nature. **b)** CATH domain diversity generated throughout evolution of ProteinGAN. At every 1200 learning steps, 64 sequences were sampled and searched for representative CATH domains (E-value <1e-6). Inset: ProteinGAN generated novel domains that are not present in existing the MDH family (left). In contrast to ProteinGAN-generated sequences, randomly introducing mutations does not expand MDH sequence diversity (Fisher’s exact test *p*-value < 8.2e-16), but rather decreases CATH domain diversity (right). For random controls, 10 000 mutated natural sequences were simultaneously searched for presence of CATH domains (see Methods). **c)** Comparison of sequence diversity between generated sequences and the natural MDH dataset. By varying sequence identity cutoffs, generated sequences group into up to 4 times more clusters than natural sequences demonstrating expanded sequence diversity. Inset shows the ratio of number of clusters (Y-axis) at different sequence identity cutoffs (X-axis). **d)** MDH activity measured by fluorescently monitoring NADH consumption using protein expression method 1 (Methods). **e)** Catalytic activity confirmed using LC-MS/MS operating in selected reaction monitoring mode for enzymes expressed using protein expression method 1 (Methods). **f)** Oxaloacetate to malate conversion yields are comparable to natural MDH enzymes as determined using mass spectrometry.

First, we evaluated ProteinGAN’s ability to capture evolutionary sequence properties, i.e. the positional amino acid variation in generated and natural sequences. Shannon entropies were computed for each position in multiple sequence alignments of the generated and natural MDH sequences (Figure 1c). The positional variability in generated sequences was highly similar to that in natural sequences, with peaks (high entropy) and valleys (low entropy) appearing at similar positions in the sequence alignment. Indeed, we observed a high correlation (Pearson’s *r* = 0.89, *p*-value < 1e-16) between the entropy values of generated and natural sequences. At conserved positions the generated sequences preserved key substrate-binding and catalytic residues (Figure 1d). Further comparative analysis of generated and natural sequences showed that even in highly variable sequence regions, the frequencies of individual amino acids were perfectly correlated (Pearson’s *r* = 0.96, *p*-value < 1e-16, Supplementary Figure 5). Moreover, for each individual sequence, ProteinGAN inferred the specific physicochemical amino acid signatures present in this enzyme class. For instance, despite high sequence diversity among generated sequences, the fractions of hydrophobic, aromatic, charged and cysteine-containing residues were the same (Wilcoxon rank sum test *p*-value > 0.05) as in natural ones. Apart from the differences in hydrophilic and polar uncharged residues (*p*-value = 7e-5 and 1e-28, respectively), the network had learned the overall amino acid composition corresponding to both the evolutionary and physicochemical constraints (Figure 1c,d; Supplementary Table 1; Supplementary Figure 6, 7).

In proteins, amino acid pairs that are remote on the primary sequence are often spatially close and interact in the 3D structure, ensuring the appropriate protein stability and function ^31^. We therefore assessed whether ProteinGAN was able to learn such local and global amino acid relationships by looking for pairwise amino acid relationships across the full length of the MDH sequences. To investigate local pairwise relationships we calculated the amino acid association measures for natural and generated sequences using the minimal proximity function Z_m_ (Santoni et al. 2016). The function Z_m_(A,B) counts, for each pairwise combination of the 20 amino acids, the average distance between amino acid A to the next amino acid B occuring in the sequence. The calculated distances can be expressed as a matrix of all pairwise combinations (Figure 1e). The matrices for the natural and generated sequences were 88% similar, showing that the amino acid positional order had been learned by ProteinGAN, capturing the local amino acid relationships existing in natural sequences. One of the main differences between the two sequence sets was in tryptophan (Figure 1e, W column), likely resulting from the fact that 22% of the natural MDH sequences used did not have tryptophan. To investigate the global amino acid relationships, we calculated the pairwise amino acid frequency distributions for all combinations of position pairs in all sequences in multiple sequence alignments. These frequency distributions were then used to calculate correlations between the training and generated sequences. Overall, we found strong correlations between the natural and generated sequences (averaged Pearson’s *r* = 0.95, Supplementary Figure 8), which demonstrated that the pairwise relationships were highly similar in both sets of sequences. To expand on this, we inspected whether the generated MDH sequences possessed the two main Pfam (Finn et al. 2014) domains “Ldh_1_N” and “Ldh_1_C” that were identified (*E*-value < 1e-10) in the natural MDH sequences. Indeed, we found that 98% of the generated sequences contained both signatures, with the rest containing one of the two domains. Collectively, our results show that ProteinGAN-generated sequences are of high quality and closely mimic natural MDH proteins, both in terms of amino acid distributions at individual sites, as well as in terms of local and long-distance relationships between pairs of amino acids present throughout the primary sequence of the MDH family.

We then explored whether ProteinGAN was able to generalize the entire protein family beyond the training dataset, i.e. to generate novel sequence diversity. Visualization of the sequence diversity of generated and natural sequences using t-distributed stochastic neighbour embedding (t-SNE) dimensionality reduction^32^ showed that a majority of natural MDH sequences grouped into large clusters (Figure 2a), as they were highly similar (median pairwise identity 92%, Supplementary Figure 9). In contrast, the generated sequences grouped into smaller clusters interpolating between the natural sequence clusters and resembled a learned manifold of the MDH sequence space (Figure 2a). To assess whether the diversity in generated sequences would contain novel and functionally relevant biological properties, we performed a search of CATH ^33^ sequence models corresponding to all known 3D structural protein domains (Methods). We first evaluated whether the network generated new structural domain diversity over the training period (Figure 2b). While the number of identified structural domains plateaued at the early stage of training (after approx. 0.2M steps), corresponding to 79% of all identified domains, additional structural CATH domains were discovered throughout the entire training process. In total, 119 novel structural sequence motifs were identified (*E*-value < 1e-6) in the generated sequences that do not exist in the training MDH dataset (Figure 2b inset), demonstrating the network’s ability to generate novel biologically relevant sequences. We next evaluated whether the generated structural domain diversity was due to chance. To test this, as a control, we randomly introduced amino acid substitutions into the natural MDH sequences, whilst preserving natural amino acid frequency distribution and the rate of mutations to mimic the natural sequence variability (Figure 2b inset, Methods). The total structural domain diversity was reduced by 38.9% in mutated natural sequences, of which 97.4% were present in natural sequences, demonstrating that random mutations did not produce biologically relevant sequence diversity (Figure 2b inset, Fisher’s exact test *p*-value < 8.2e-16). Using clustering analysis based on sequence similarity we observed that on average over 95% of the generated sequences were not more than 10% similar to each other (90% sequence identity within the cluster, Figure 2c), in contrast to only 17% of the natural sequences at the same sequence identity level. This shows that the generated sequences expanded the currently known sequence space of the MDH family up to 4-fold (Figure 2c inset).

Finally, considering that random amino acid substitutions typically result in a decline or even complete loss of protein activity ^4,10–16^, we experimentally tested whether the ProteinGAN-generated MDH sequences were catalytically active *in vitro*. We selected 60 representative generated sequences within a range of 45% to 98% pairwise sequence identity to natural MDH and with 7 to 157 amino acid substitutions compared to their closest MDH neighbour (Supplementary Figure 10, Supplementary Table 2), of which 55 were successfully synthesized and cloned into an expression vector. Production of recombinant proteins in *Escherichia coli* and purification using affinity chromatography yielded 11 protein variants (Method 1) that could be purified from the cell lysate soluble fraction (Supplementary Table 3, Supplementary Figure 11). With the aim to identify additional soluble proteins, we repeated the experiment (Method 2) under growth conditions favouring protein folding and solubility using ArcticExpress *E.coli* strain, expanding the number of purified soluble proteins to a total of 19 (Supplementary Table 3, 35% of all synthesized protein variants). This is comparable to other systematic studies, which typically obtain soluble protein for 20% to 40% of all tested constructs ^34–36^. The purified proteins were assessed for MDH activity (Supplementary Figure 2) by monitoring NADH consumption using a spectrophotometer. 13 of the 19 soluble enzymes, including a variant with 106 amino acid substitutions (66% identity to the closest existing enzyme, Supplementary Figure 12), showed MDH catalytic activity (Figure 2d, Supplementary Table 3, Supplementary Figures 13, 14). Furthermore, for the subset of 8 purified enzymes for which the protein amount could be accurately quantified the generated MDH proteins displayed similar reaction rates as wild-type enzymes (Supplementary Figure 13). These enzymes were also confirmed, by LC-MS/MS, to convert oxaloacetate to malate with reaction yields comparable to commercial MDH enzyme controls (Figure 2e, f).

In conclusion, here we present a generative adversarial network, ProteinGAN, that successfully captures the natural properties of proteins and enables discovery of novel functional sequences. *In vitro* experiments confirm that a large portion (24%) of the generated enzymes are soluble and many display catalytic activities comparable to - or surpassing that of - natural enzymes (Figure 2d,e,f Supplementary Figure 11,13). The generated functional enzymes contain up to 106 mutations compared to the closest natural malate dehydrogenase (Supplementary Figure 12), while retaining functionally relevant sequence motifs and the correct position-specific amino acid composition that preserves long-range amino acid interactions (Figure 1). Since ProteinGAN enables large leaps to unexplored sections of the functional sequence space (Figure 2a), it opens up the biochemical exploration of the highly diverse enzymes populating this space. Such enzymes may have catalytic properties that differ significantly from those found in natural enzymes, as they have not evolved with the constraint to carry out specific functions in living organisms as natural enzymes have. Navigating the fitness landscape to find even one such enzyme using current methods, including random mutagenesis ^4^ and recombination of homologous proteins ^8,9^, would be highly laborious or not even feasible. Due to the exponential decline in protein fitness with the number of random mutations ^11,37^ or the number of parent molecules that are used to generate the recombination libraries ^38,39^, current methods are fundamentally limited in the sequence space that can be explored by their use. The expanded functional sequence diversity provided by ProteinGAN (Figure 2b,c) may also provide suitable, non-natural, starting points for protein engineering ^40^, with great potential for applications in biocatalysis ^41^. We speculate that further development of the ProteinGAN framework will enable even greater leaps in sequence space and may also make the method applicable to smaller enzyme families. Future work should explore whether ProteinGAN can be trained on entire protein families where, despite sharing sequence similarity, its members perform distinct functions.

## Methods and materials

### Neural network architecture details

The GAN architecture consisted of two networks - a discriminator and a generator - each of which used ResNet blocks ^42^ (Supplementary Figure 1). Each block in the discriminator contained 3 convolution layers with filter size of 3×3, 2 batch normalization layers ^43^ and leaky ReLU activations ^44^. The generator residual blocks consisted of two transposed convolution layers, one convolution layer with the same filter size of 3×3 and leaky ReLU activations. Each network had one self-attention layer ^30^. Transposed convolution technique was chosen for up-sampling, as it yielded the best results experimentally (Supplementary Figure 15). For loss, non-saturating loss with R1 regularization ^45^ was used (Supplementary Figure 16). To ensure training stability, Spectral normalization ^46^ was implemented in all layers.

The input to the discriminator was one-hot encoded with a vocabulary size of 21 (20 canonical amino acids and a sign that denoted a space at the beginning or end of the sequence). The generator input was a vector of 128 values that were drawn from a random distribution with mean 0 and standard deviation of 0.5, with the exception that values whose magnitude was more than 2 standard deviations away from the mean were re-sampled. The dimensions of generated outputs were 512×21, wherein some of the positions denoted spaces. The implementation of ProteinGAN can be accessed at https://github.com/biomatterdesigns/ProteinGAN.

### Network training data

Bacterial malate dehydrogenase (MDH) sequences were downloaded from Uniprot on January 10th 2019 ^47^. Sequences longer than 512 amino acids or containing non-canonical amino acids were filtered out. The final dataset consisted of 16 898 sequences, which were clustered into 70% identity clusters using the MMseq2 tool ^48^ for balancing the dataset during the training process. 20% of the clusters with less than 3 sequences were randomly selected for validation (192 sequences) and the rest of the dataset was used for training (16 706 sequences).

### Network training process

The ratio 1:1 between generator and discriminator training steps was selected (Supplementary materials, Supplementary Figure 17). ADAM algorithm ^49^ was used to optimize both networks. Throughout the training, the learning rate was gradually decreased from 1e-3 to 5e-5 for both the generator and the discriminator. To avoid bias towards sequences with large number of homologues, smaller clusters were dynamically up-sampled during the training. In order to track the performance, along with GAN losses, generated data was constantly evaluated. Without halting the training process, every 1200 steps generated sequences were automatically aligned with the training and validation datasets using BLAST ^50,51^. Throughout the training, BLOSUM45, *E*-value and identity scores as well as standard deviation of the discriminator layer were calculated and monitored (Supplementary Figure 3, 18, 19, 20). ProteinGAN was trained for 2.5 M steps with a batch size of 64 (one step consisted of 64 sequences). The training took 210 hours (~9 days) on NVIDIA Tesla P100 (16 GB).

### Generated sequences bioinformatic analysis

#### Multiple sequence alignments

Multiple sequence alignments (MSA) (Figure 1) were calculated using using Clustal Omega ^52^ by merging natural and generated datasets in equal amounts. To calculate further Shanon entropies, after MSA we split the alignment corresponding to generated and training sequences. Columns having more than 75% of gaps in either dataset were removed from further analysis. For each column in MSA, Shannon entropy was calculated as follows: 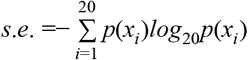, where *p*(*x*_*i*_) is the frequency of amino acid *i* occurring at a column of MSA.

#### Amino acid pair association matrices

Amino acid pair association matrices were calculated for every possible pair in a sequence and averaged over the whole dataset. The association score was used as reported in the original article (Santoni et al. 2016), where Z_m_ is expressed as follows: 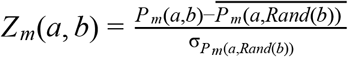. Here 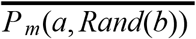 and 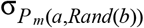 are the average and the standard deviation of the randomly shuffled sequence association score for the same pair. The association function for scoring was selected as the minimal proximity function: 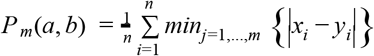. Here for each position *x*_*i*_ of amino acid *a*, the closest occurrence of amino acid *b* at position *y*_*i*_ is identified and the average of the distances between the pairs is calculated. In our implementation, if a sequence does not contain a certain amino acid, Null value is returned for the pairs containing the amino acid.

#### Sequence clustering

Sequence clustering was performed using MMseqs2 ^48^ with easy-cluster option and required sequence identity cutoff.

#### Pfam/CATH domain search

All sequences generated by ProteinGAN were classified using HMMER3 ^53^ search over Pfam 32.0 database ^54^. HMMs for each CATH representative domain from the sequence clusters at 35% sequence identity (v4.1) were downloaded from CATH database repository ^33^. To avoid biases in sequence scoring, generated sequences together with the natural sequences in equal quantities were appended to the same file and in all tests were searched simultaneously using hmmsearch tool with default options ^53^. To evaluate whether generated domain diversity was not due to chance, we chose a random subset of 10 000 natural sequences and mutated them by randomly introducing on average 100± 30 sd substitutions of amino acids (corresponding to median identity of generated sequences Figure 1b) that were uniformly sampled of natural amino acid probability distribution. Generated, natural and mutated sequences (10 000 of each) were searched as one database and hits were considered significant with E-value < 1e-6. The analyzed sequences were generated by latest checkpoint model (~2.5M training steps).

#### t-SNE plot generation

A distance matrix of cluster representatives was used as the t-SNE input. To get cluster representatives, first, the number of sequences in both datasets were equalized by taking 13,272 sequences from natural and generated datasets. These sequences were independently clustered using MMseqs2 ^48^ with 70% minimal sequence identity. This generated 926 clusters of natural sequences and 3,778 clusters of generated sequences. Representative sequences of these clusters were chosen based on MMseqs2 output. From the representative sequences a distance matrix was generated using Clustal Omega ^52,55^. The distance matrix was used with the scikit-learn t-SNE module ^56^ with default settings (early exaggeration 12, learning rate 200, maximum number of iterations: 1000) except that the embedding generation perplexity was set to 7. Coordinates given by t-SNE were used for plotting, the size of a given dot was visualized based on the cluster size it represents.

#### Visualization of GAN training

Sequences generated during the training period were sampled at 14 different times. For each of the 14 checkpoints, 64 sequences were taken, every checkpoint was taken after *i*(*x*) = 2^*x*^ ∗ 300 (where x is number of checkpoints) GAN steps, with the first checkpoint replaced with 1 instead of 300. For all the generated sequences, a global identity to the closest sequence in training dataset was calculated. Identities of each checkpoint were plotted.

#### Correlation of distant dimer pairs

To calculate the correlation of close and distant dimer pairs between the datasets, the total number of individual dimers for every possible MSA position pair was calculated: *ds* = {*m*_*a*1,*a*1_, *m*_*a*1,*a*2_, …, *m*_*a*21,*a*21_}, here *m*_*ai*,*aj*_ was a set of dimer *a*_*i*_*a*_*j*_ counts over each position pair of a multiple sequence alignment (*m*_*ai*,*aj*_ = {*d*_1,1_, *d*_1,2_, …, *d*_*n*,*n*_}). Each number of *m*_*ai*,*aj*_ set was calculated by summing *a*_*i*_*a*_*j*_ dimers over all sequences of MSA in positions *n* and *m*: 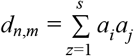 here *s* is the total number of sequences in MSA. *ds* was calculated for each dataset and for each member of the *ds* set, Pearson’s r was calculated between the datasets (natural and generated). These correlations were plotted as a heatmap (Supplementary Figure 8). For *ds* calculation only columns containing less than 75% of gaps in both natural or generated datasets were used.

### Experimental validation of generated enzymes

The sequences generated by ProteinGAN were synthesized, cloned into the pET21a expression vector and sequence-verified by Twist Bioscience. In addition to the enzyme sequence a C-terminal linker and four histidines (AAALEHHHH) were added, resulting in a deca-His-tag in the final construct (which includes six histidines derived from the expression vector), to enable downstream affinity purification. Method 1: The constructs were transformed into the BL21(DE3) *E. coli* expression strain. From the resulting transformation mixture 15 μl was used to inoculate 500 μl LB broth supplemented with 100 μg/ml carbenicillin. Cells were grown overnight at 32°C in a 96 deep well plate with 700 rpm orbital shaking. Protein expression was achieved by diluting the overnight cultures 1:30 into 1 ml autoinduction TB including trace elements (Formedium, UK) supplemented with 100 μg/ml carbenicillin and grown for 4 h at 37°C, followed by overnight growth at 18°C and 700 rpm shaking. Cells were collected by centrifugation and the cell pellets frozen at −80°C overnight. To purify the recombinant proteins, cells were thawed, resuspended in 200 μl lysis buffer (50 mM HEPES pH 7.4, 5% glycerol, 300 mM NaCl, 0.5 mM TCEP, 0.5 mg/ml lysozyme, 10 U/ml DNaseI, 2 mM MgCl_2_), and incubated for 30 min at room temperature. To improve lysis triton-X-100 was added to a final concentration of 0.125% (v/v), and the cells were frozen in −80°C for 30 min. After thawing in a room temperature water bath, the lysates were spun down for 10 min at 3000 × g to remove cell debris, and the supernatants were transferred to a new 96-well plate with 50 μl Talon resin in each well (Takara Bio, Japan). Unspecific binding of proteins to the resin was reduced by adding imidazole to a final concentration of 10 mM in each well. The plate was incubated at room temperature for 30 min with 400 rpm shaking, after which the lysates with the beads were transferred to a 96-well filter plate (Thermo Scientific, USA, Nunc 96-well filter plates), placed over a 96-well collection plate, and centrifuged for 1 min at 500 x g in a swing-out centrifuge. The resin was washed three times with 200 μl wash buffer (50 mM HEPES pH 7.4, 5% glycerol, 300 mM NaCl, 0.5 mM TCEP, 40 mM imidazole), and the proteins were eluted from the resin in two 50 μl fractions using elution buffer (50 mM HEPES pH 7.4, 5% glycerol, 300 mM NaCl, 0.5 mM TCEP, 250 mM imidazole). The two eluate fractions were combined and transferred to a 96-well desalting plate (Thermo Scientific, USA, Zeba Spin Desalting Plate, 7K MWCO) pre-equilibrated with sample buffer (50 mM HEPES pH 7.4, 5% glycerol, 300 mM NaCl, 0.5 mM TCEP). The plate was spun down 1000 x g for 1 min, and collected proteins were analysed by SDS-PAGE followed by Coomassie staining. The soluble proteins were carried on for further characterisation.

To test for malate dehydrogenase activity, an aliquot of purified protein was added to a reaction mixture containing 0.15 mM NADH, 0.2 mM oxaloacetic acid and 20 mM HEPES buffer (pH 7.4). The final reaction volume was 100 μl and the reaction was carried out at room temperature in a UV-transparent 96-well half-area plate (UV-Star Microplate, Greiner, Austria). Activity was measured in triplicates by following NADH oxidation to NAD+, with absorbance reading at 340 nm performed every 30 sec for 15 min in a BMG Labtech SPECTROstar Nano spectrophotometer. Un-specific oxidation of NADH was monitored in no-substrate controls, and these values were subtracted from the other samples. Conversion from absorption values to NADH concentration was carried out using an extinction coefficient of 6.22 mM. For calculation of kinetic parameters, 10 nM of each protein was assayed with a range of oxaloacetate concentrations.

LC-MS/MS quantification was performed for selected active enzymes. The activity assay was performed as outlined above, in triplicates, with protein concentrations ranging between 10 and 250 nM. Reactions were terminated after 45 min by diluting the assay mixtures in water to 1 μg/ml starting concentration of oxaloacetate. For chromatographic separation a Zorbax Eclipse Plus C18 50 mm × 2.1 mm × 1.8 μm (Agilent) with an Nexera series HPLC (Shimadzu) was used. Mobile phase A was composed of H20 (MiliQ HPLC grade) with 0.1% Formic acid (Sigma); mobile phase B was Methanol (Sigma) with 0.1% Formic acid (Sigma). The oven temperature was 40°C. The chromatographic gradient was set to consecutively increase from 0% to 100%, hold, decrease from 100% to 0% and hold, in 60 sec, 30 sec, 30 sec and 30 sec, respectively. The autosampler temperature was 15°C and the injection volume was 0.5 μL with full loop injection. For MS quantification a QTRAP® 6500 System (Sciex) was used, operating in negative mode with Multiple Reaction Monitoring (MRM) parameters optimized for Malic acid based on published parameters ^57^. Electrospray ionization parameters were optimized for 0.8mL/min flow rate, and were as follows: electrospray voltage of −4500 V, temperature of 500 °C, curtain gas of 40, CAD gas set to Medium, and gas 1 and 2 of 50 and 50 psi, respectively. The instrument was mass calibrated with a mixture of polypropylene glycol (PPG) standards. The software Analyst 1.7 (Sciex) and MultiQuant 3 (Sciex) was used for analysis and quantitation of results, respectively.

Additionally (Method 2), to obtain additional soluble proteins, MDH constructs were transformed into ArcticExpress competent cells (Agilent technologies, USA). The transformants were inoculated into 500 μL of LB media with 15 μg/mL of Gentamicin and 50 μg/mL of Ampicillin and grown overnight at 30° C in a Thermomixer Comfort Eppendorf thermomixer (Eppendorf, Germany). 250 μL of overnight culture were transferred to 10 mL (dilution 1:40) of semi-synthetic media (1 % Tryptone, 0.5% Yeast extract, 0.268 % (NH4)2SO4, 0.15% NH4Cl, 0.6 % KH2PO4, 0.4% K2HPO4, 1% Glycerol, pH 7.0) supplemented with 15 μg/mL of Gentamicin and 50 μg/mL of Ampicillin. Cells were cultivated at 37 ° C for 2 hours, until OD600 reached 0.6-0.8, then the media were enriched with 0.5 M of saccharose. Induction was carried out at 12°C with 0.5 mM IPTG overnight. The cells were harvested by centrifugation (4000 x g 10 min at 4° C), resuspended in 0.1 M potassium phosphate buffer, pH 7.0 and then sonicated on ice in 2.0 mL tubes at 30% amplitude for 5 min of total ON time (30 s on/30 s off) by using the Bandelin SonoPuls HD 2070 homogeniser (BANDELIN, Germany).

To remove cell debris, the lysates were centrifuged at 16 000 x g, 4° C. The soluble recombinant MDH mutants were purified by using HisPur™ Ni-NTA Spin columns (ThermoFisher Scientific, USA). The columns with loaded supernatants were washed with Wash buffer (0.1 M Potassium phosphate buffer, pH7.4, NaCl 250 mM and 40 mM imidazole). Proteins were eluted with Elution buffer (0.1 M Potassium phosphate buffer, pH 7.4, NaCl 250 mM and 300 mM). The eluted fractions were dialysed against 0.1M Potassium phosphate buffer, pH 7.4. The concentration of the proteins was determined using NanoDrop 2000 (Thermofisher Scientific, USA). The aliquots of total lysate, soluble lysate fraction and purified protein were loaded onto SDS PAGE 15%.

The malate dehydrogenase activity was measured at 25 ° C in 96-well flat bottom UV transparent plate (UV-Star Microplate) (Greiner Bio-One, Austria). The reaction mixture (final volume 200 μL) contained an aliquot of purified protein, freshly prepared 0.15 mM NADH and 0.2 mM oxaloacetic acid, and 0.1 M Potassium phosphate buffer, pH 7.4. The absorbance reading was done at 340 nm every 5 seconds for 3 minutes in a BioTek PowerWave XS microplate reader (Biotek, USA). For NADH at 340 nm 6.22 mM cm-1 extinction coefficient (εM) was used. The path length (l) in the microplate was calculated according A=c × εM × l.

## Acknowledgements

We thank Greta Stonyte, Juozas Nainys, Simran Aulakh and Clara Correia-Melo for comments on the manuscript. L.K. and R.M. were supported by the Agency for Science, Innovation and Technology (Lithuania) grants No. 31V-59/(1.78)SU-1687 and 01.2.2-MITA-k-702-04-0001. A.Z. is a SciLifeLab fellow.

