## Supplementary Information for "Expanding functional protein sequence space using generative adversarial networks"

1 - Biomatter Designs, Sauletekio al. 7, LT-10257, Vilnius, Lithuania

2 - Institute of Biotechnology, Life Sciences Center, Vilnius University, Saulėtekio al. 7, LT-10257, Vilnius, Lithuania

3 - Department of Biology and Biological Engineering, Chalmers University of Technology, Kemivägen 10, SE-412 96, Gothenburg, Sweden

4 - Institute of Biochemistry, Life Sciences Center, Vilnius University, Saulėtekio al. 7, LT-10257, Vilnius, Lithuania

5 - Chalmers Mass Spectrometry Infrastructure, Chalmers University of Technology, Kemivägen 10, SE-412 96, Gothenburg, Sweden

6 - Science for Life Laboratory, Tomtebodavägen 23a, SE-171 65, Stockholm, Sweden

*Equal contribution

### Table of contents

Supplementary Methods … p.2 - p.4

Supplementary Figures … p.5 - p.29

Supplementary Tables … p.30 - p.41

Supplementary References … p.42 - p.43

### Methods

#### Generative Adversarial Networks

##### Mode collapse

A common issue with GAN architectures is mode collapse [1], which manifests itself when the generator generates a limited diversity of outputs regardless of the inputs. In order to overcome the mode collapse issue we used the Mini Batch Discriminator (MBD) approach [2]. MBD works as an extra layer in the network that computes the standard deviation across the batch of examples (batch contains only real or only fake sequences). If the batch contains a small variety of examples, the standard deviation will be low and the discriminator will be able to use this information to lower the final score for each example in the batch. Moreover, to further reduce mode collapse occurrence we balanced the sampling frequency of the training dataset clusters.

##### Dilation

Convolution filters are very good at detecting local features, but they have limitations when it comes to long distance relationships [3]. As a result, a large portion of deep learning algorithms use RNNs (Recurrent Neural Network) when it comes to sequences. However, it has been shown that convolution filters with dilation outperform RNNs [3]. The idea of dilation is to increase the receptive field without increasing the number of parameters, by introducing gaps into convolution kernels. Dilation rate was applied to one convolution filter in each residual block (Supplementary Figure 21). The dilation rate was doubled in each consecutive block. In this way, by the last layer of the network, filters had a large enough receptive field to learn long-distance relationships.

##### Self-Attention

Different areas of a protein have different responsibilities in overall protein behaviour. In order for the network to capture this, self-attention mechanism [4] were implemented (Supplementary Figure 21). The self-attention mechanism consists of a number of layers that highlights different areas of importance across the entire sequence and allows the discriminator to check that parts in distant portions of the protein are consistent with each other.

##### Upsampling

We explored 3 widely used up-sampling techniques to choose the one that is best suited for protein generation: nearest-neighbor interpolation, transposed convolutions [5] and sub-pixel shuffle [6]. Based on experiments (Supplementary Figure 15), transposed convolutions were chosen for all upsampling layers in the final solution.

#### Loss

The key component to successful performance of neural networks is the loss function. There are a number of different loss variants for GANs that work well for image generation. We therefore implemented and evaluated a number of different losses.

We experimented with non-saturating [7], non-saturating with R1 regularization [8], hinge [9–11], hinge with relativistic average [12], Wasserstein [13] and Wasserstein with gradient penalty [14] losses. The non-saturating loss with R1 regularization was chosen for ProteinGAN due to its significant performance increase (Supplementary Figure 16).

#### Ratio between Discriminator and Generator steps

GANs commonly suffer from unreliable gradients due to poor performance of the discriminator [15]. There are two common techniques to alleviate this issue: a different number of training steps [13] and different learning rates [16]. The aim of these techniques is to have the discriminator perform better than the generator all of the time, thus achieving more reliable gradients. However, in our case, the discriminator was performing well even with a 1:1 ratio and identical learning rates (Supplementary Figure 17, 22). Furthermore, with these parameters, GAN training was more efficient. For example, training the generator for 10k steps took 1h 22min, when the ratio was 1:1, and 2h 5min, when the ratio was 1:2 (Supplementary Table 4).

#### Negative results

Even though the final solution used one-hot to encode categorical variables, we spent significant time on exploring embeddings. We experimented with self-learnt, pre-trained embeddings as well as with physicochemical properties of amino acids derived from the AAindex database [17] that served as a static embedding. Using embeddings, the generator generates physical/chemical (or in self-learnt/pre-trained case - arbitrary) properties of amino acids in each position, which later can be mapped to the closest amino acid. Unfortunately, this approach did not yield successful results. We hypothesized that the cause of this was that the input still appeared as a categorical variable despite the fact that it was encoded with continuous values. Even though introducing some randomness into real sequences alleviated the issue, it did not match the performance that was achieved by one-hot encoding.

Furthermore, we explored the idea of transfer learning. We adopted the BERT model [18] to work with amino acids instead of English words and trained it on the Uniprot database to predict masked amino acids. We used the trained model to encode the training sequences and to decode generated sequences. This approach slowed down the training and did not yield great results, and was thus abandoned.

We also experimented with a recurrent type of networks that offer the possibility of generating variable length sequences. However, because of parallelization issues of the networks, RNN type GANs were extremely slow to train and did not show promising results.

### Supplementary Figures

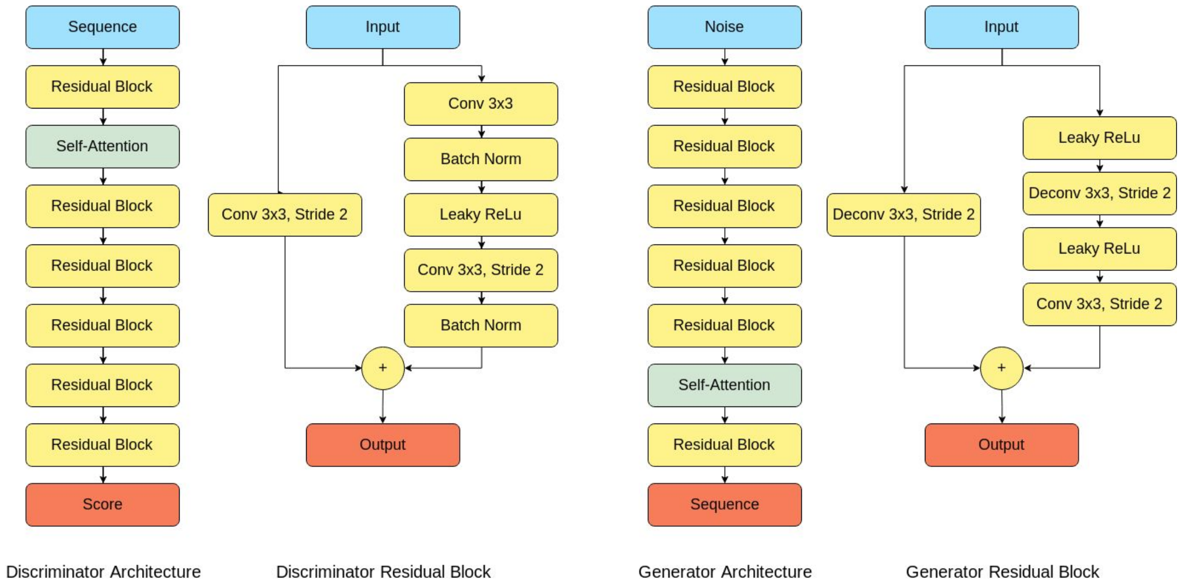

**Supplementary Figure 1 |** Graphical representation of residual blocks and high level architecture of the discriminator and generator networks.

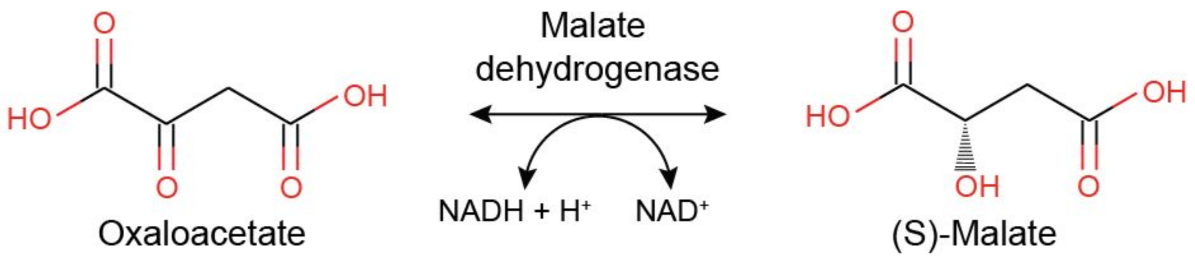

**Supplementary Figure 2 |** The reaction catalyzed by malate dehydrogenase (EC1.1.1.37). The reaction rate is determined by measuring the decrease in absorbance at 340 nm resulting from the oxidation of NADH.

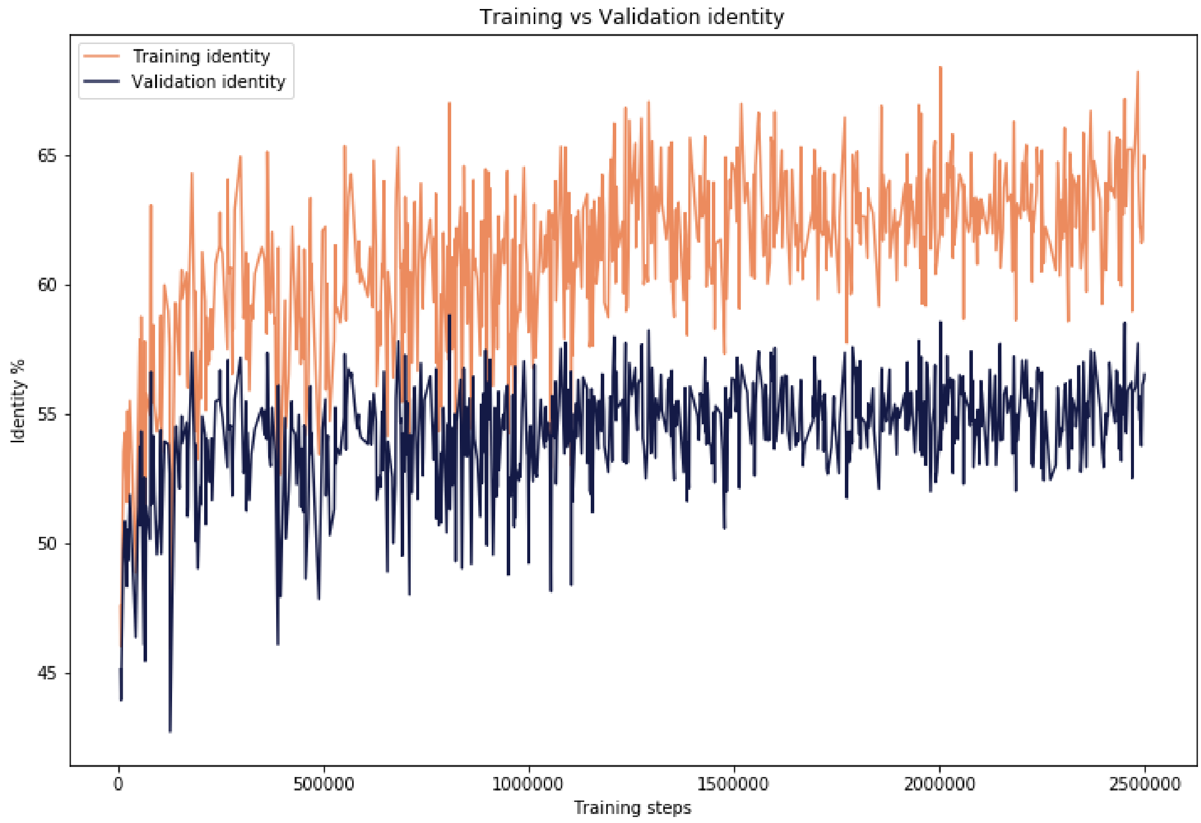

**Supplementary Figure 3 | Average identity of batches of generated sequences when compared with training and validation sequences.** Identity was calculated using BLAST tool with the BLOSUM45 matrix against sequences in the training and validation datasets separately.

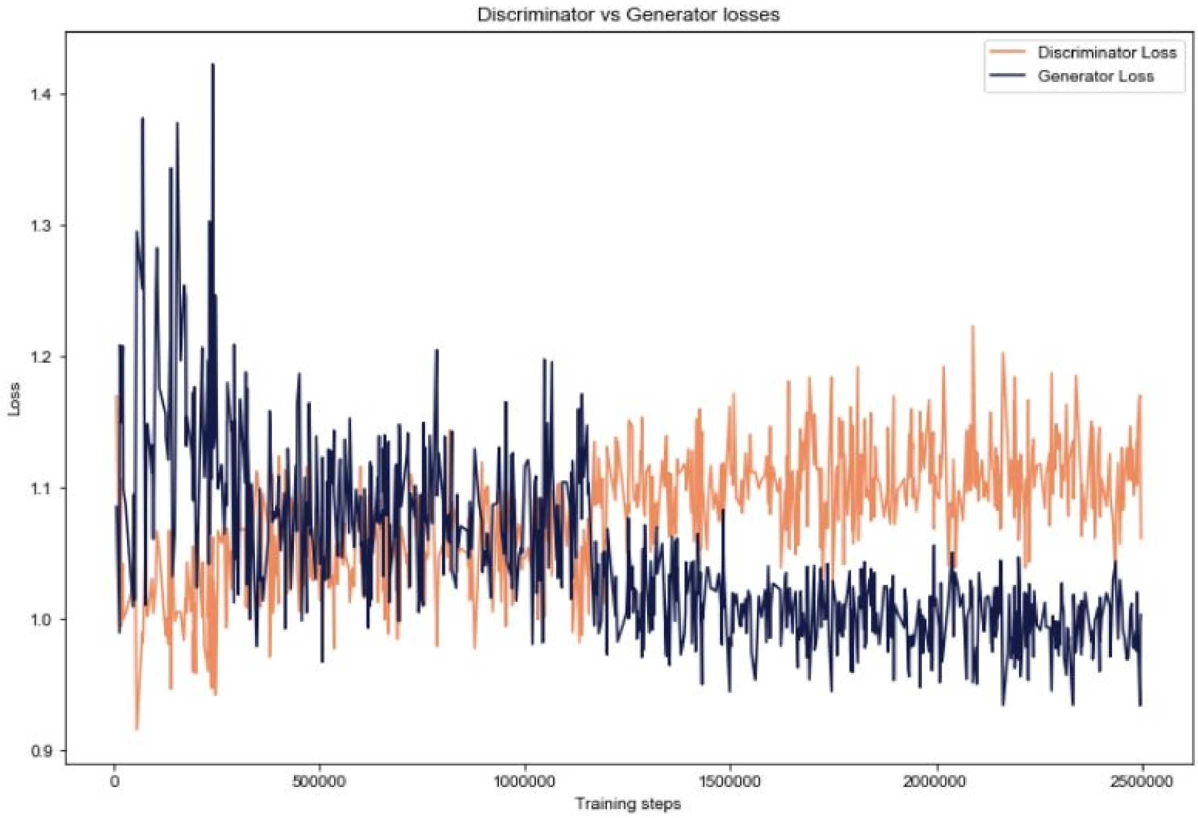

**Supplementary Figure 4 | Losses of generator and discriminator during training period.** Generator and Discriminator losses become relatively stable after the initial training phase and eventually plateaued. Note that it is impossible to determine whether networks stopped learning at this time, as the improvement of the generator could be offset by the improvement of the discriminator's ability to distinguish real from generated data.

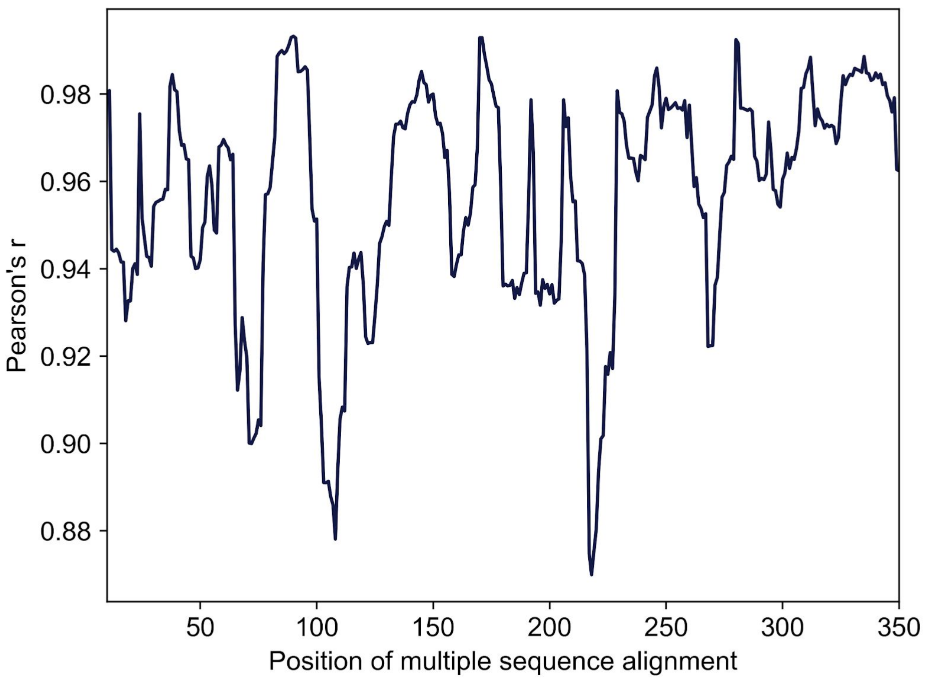

**Supplementary Figure 5 | Ability of GAN to recreate the positional amino acid distribution shown as Pearson's correlation coefficient for generated and natural sequences, estimated from multiple-sequence alignment.** Positions with lower correlation coefficients matched positions with higher sequence variability. Only positions with a number of gaps below 75% are represented, moving average of 12 positions is used.

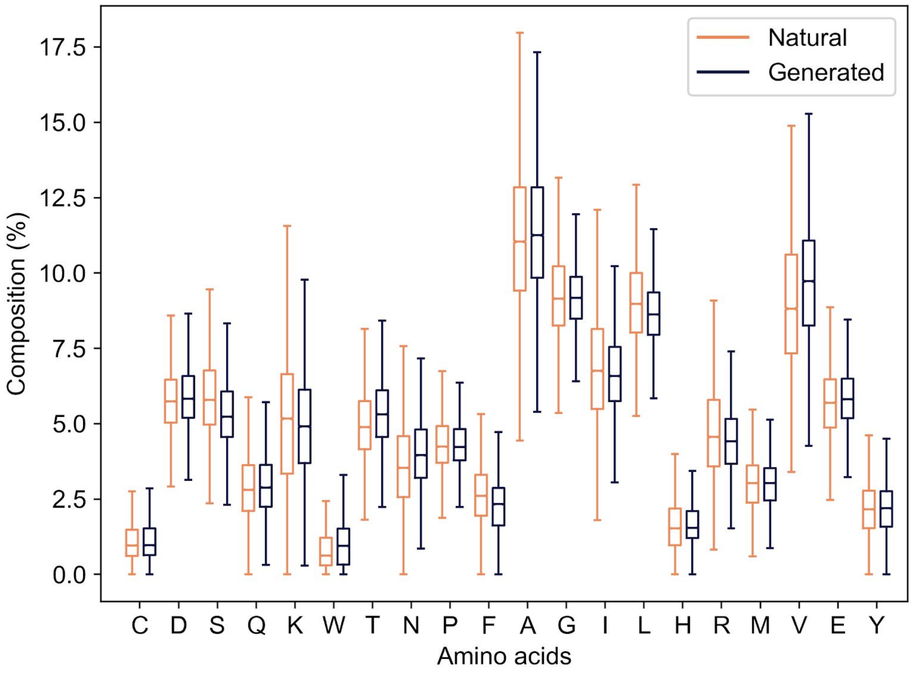

**Supplementary Figure 6 | Amino acid composition of raw sequences.** Generated sequences show a highly similar amino acid profile and compositional variability to the natural sequences.

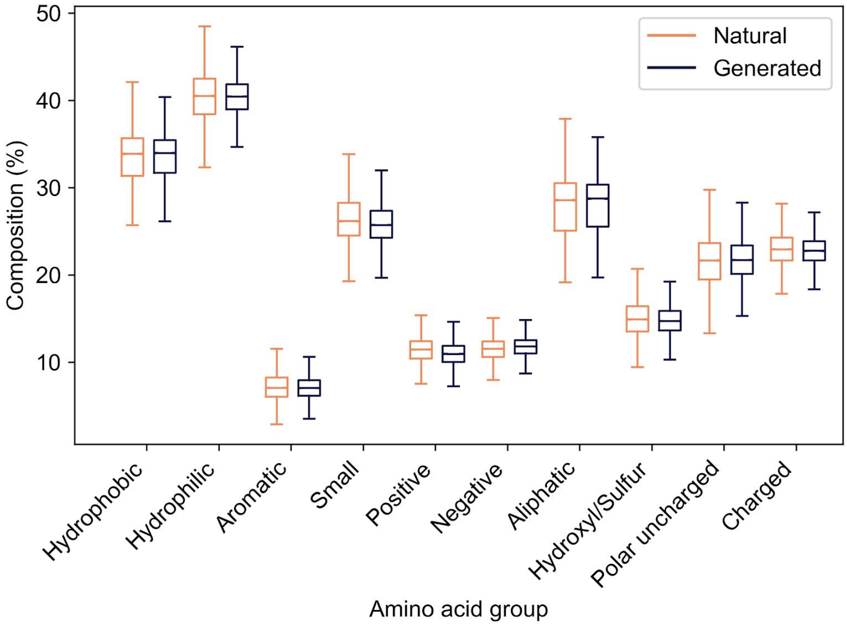

**Supplementary Figure 7 | Boxplot of percental amino acid composition of raw sequences grouped by physicochemical properties.** ProteinGAN is able to replicate the specific physicochemical properties found in natural sequences.

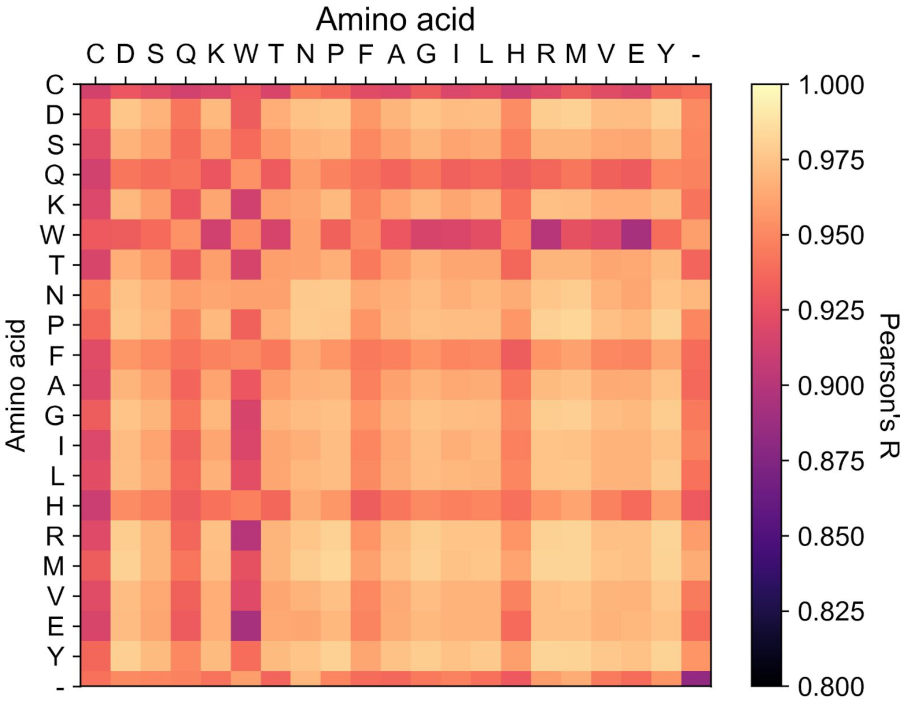

**Supplementary Figure 8 | Amino acid pair correlations of generated and natural sequences.** Every point on the map represents the correlation of amino acid pairs between two different datasets. High correlation denotes that the same pairwise long-distance amino acid interactions were found in both datasets.

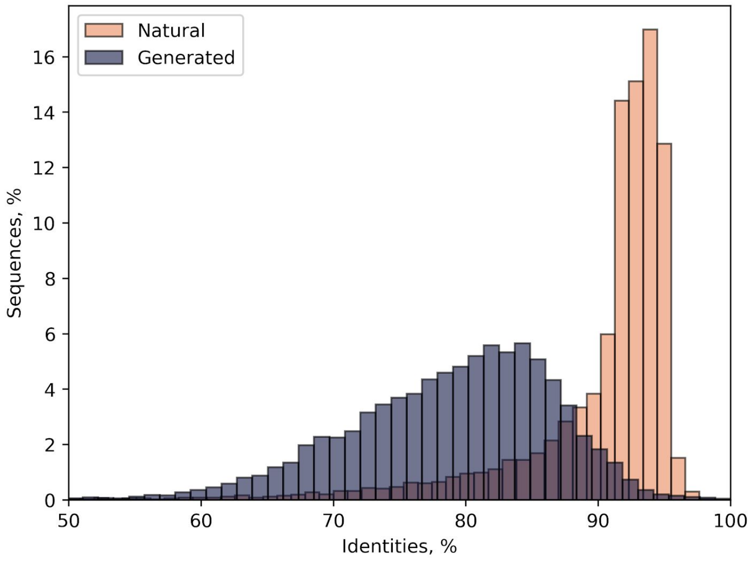

**Supplementary Figure 9 | Global identities of generated and training sequences to the closest sequence in their own dataset.** Generated sequences show a higher sequence to sequence variability than the sequences in the training dataset.

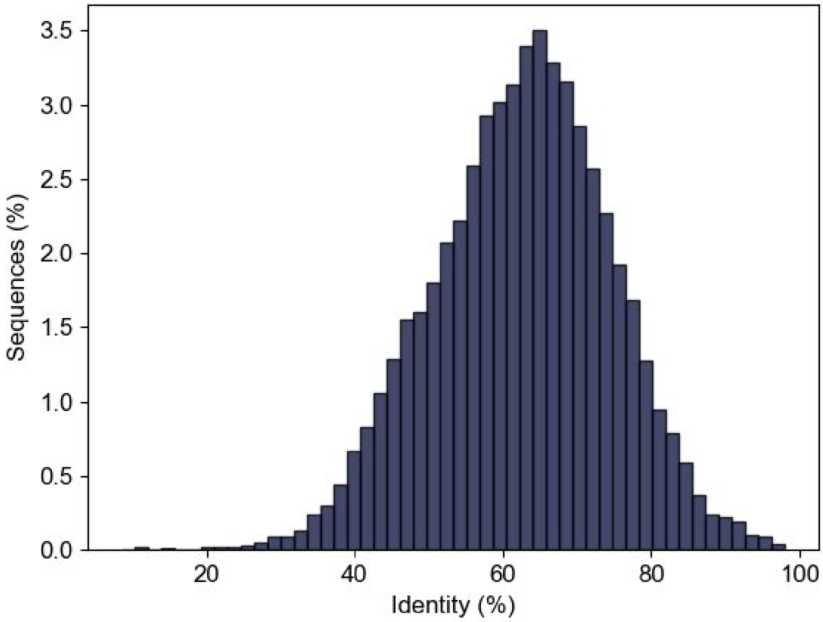

**Supplementary Figure 10 | Global pairwise sequence identity of the generated sequences to the closest sequence in the training dataset.**

**A.**

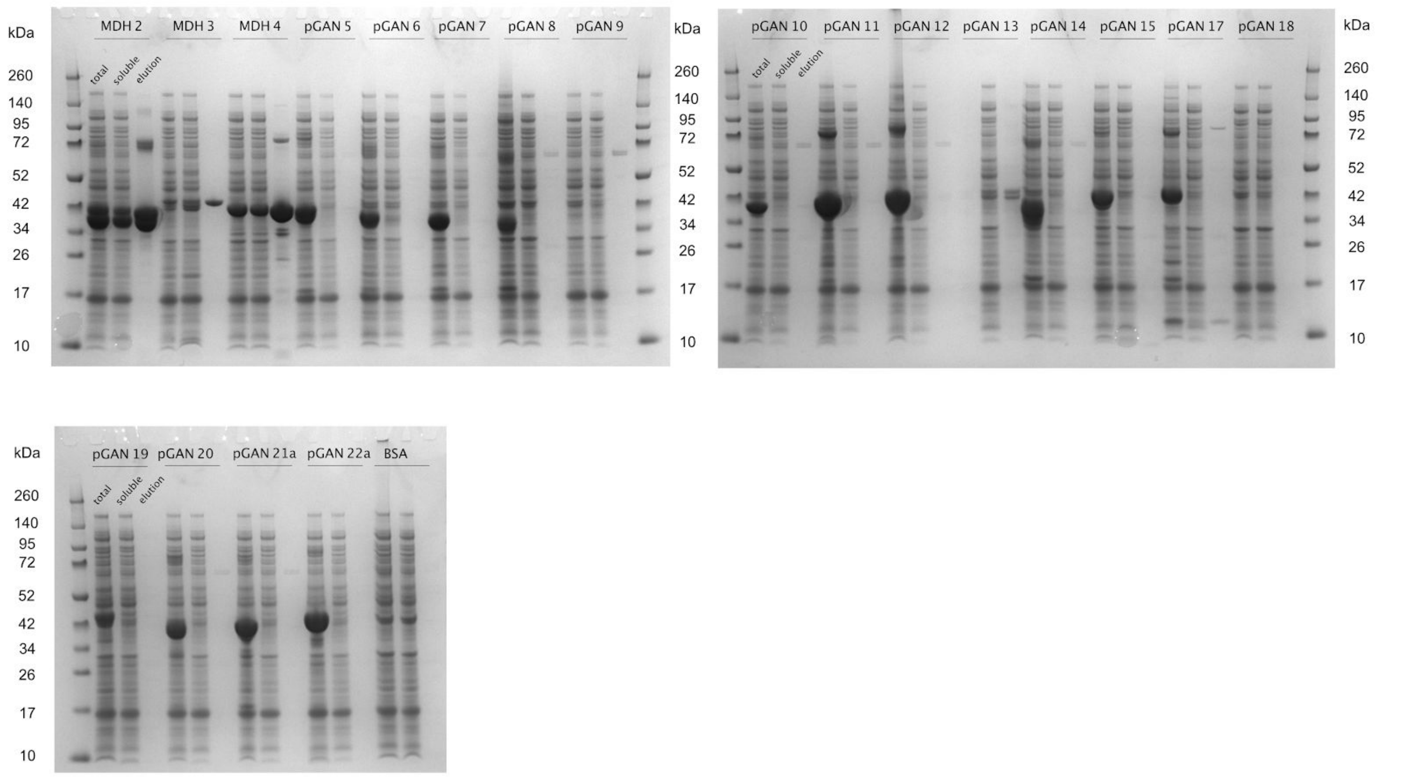
kDa
MDH 2
MDH 3
MDH 4
pGAN 5
pGAN 6
pGAN 7
pGAN 8
pGAN 9
total
soluble
elution
260
140
95
72
52
42
34
26
17
10
pGAN 10
pGAN 11
pGAN 12
pGAN 13
pGAN 14
pGAN 15
pGAN 17
pGAN 18
pGAN 19
pGAN 20
pGAN 21a
pGAN 22a
BSA

**B.**

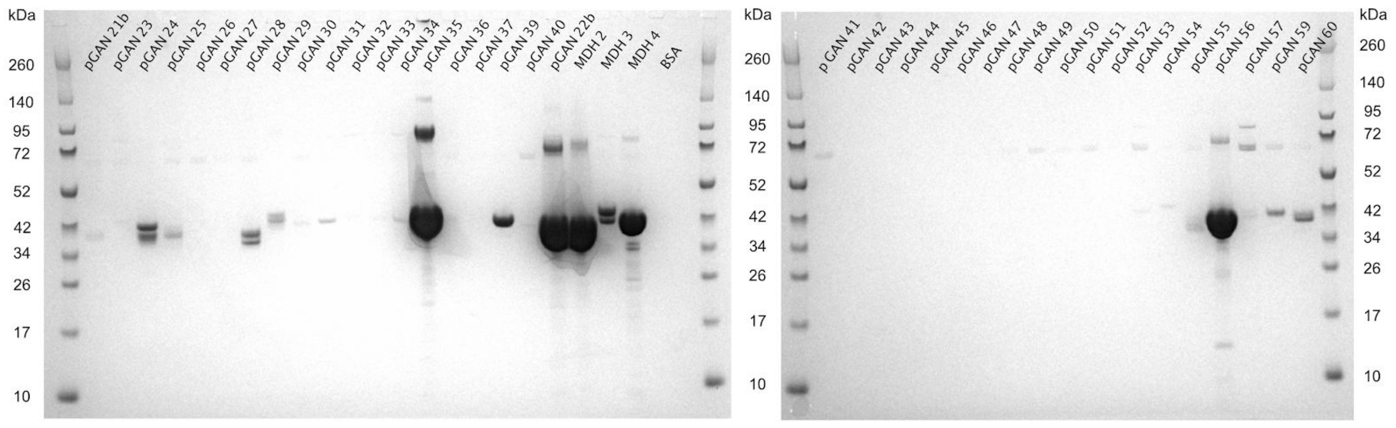
kDa
pGAN 21b
pGAN 23
pGAN 24
pGAN 25
pGAN 26
pGAN 27
pGAN 28
pGAN 29
pGAN 30
pGAN 31
pGAN 32
pGAN 33
pGAN 34
pGAN 35
pGAN 36
pGAN 37
pGAN 39
pGAN 40
pGAN 22b
MDH 2
MDH 3
MDH 4
BSA
260
140
95
72
52
42
34
26
17
10
p GAN 41
pGAN 42
pGAN 43
pGAN 44
pGAN 45
pGAN 46
pGAN 47
pGAN 48
pGAN 49
pGAN 50
pGAN 51
pGAN 52
pGAN 53
pGAN 54
pGAN 55
pGAN 56
pGAN 57
pGAN 59
pGAN 60

**C.**

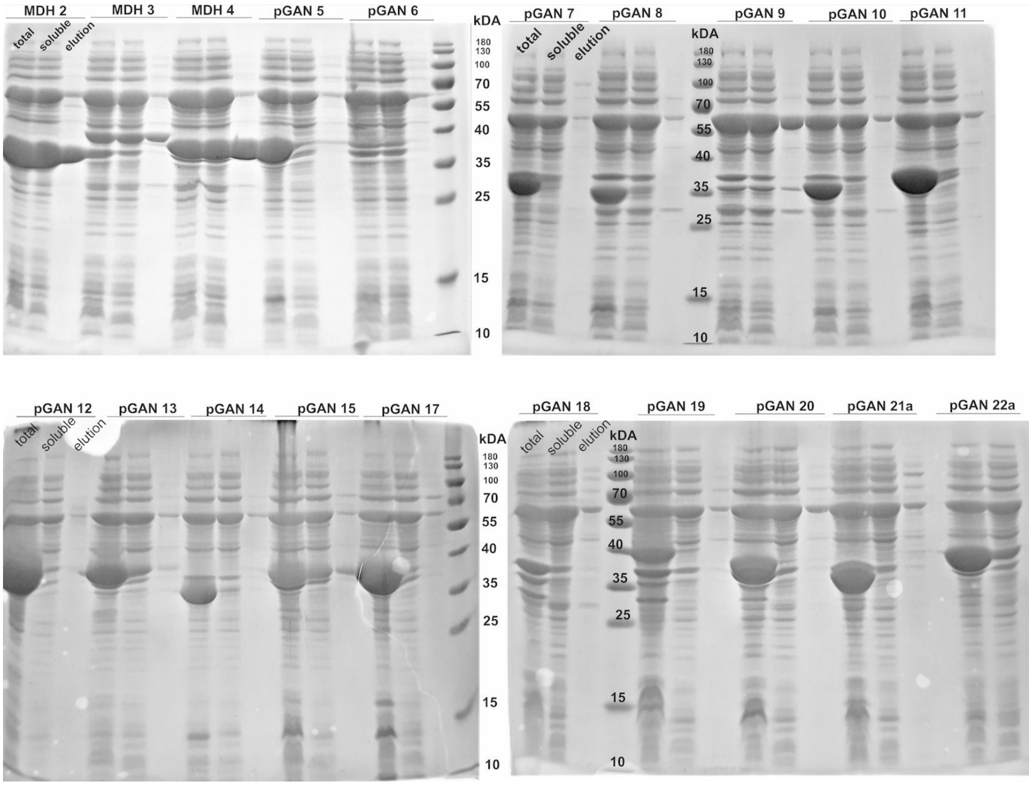
MDH 2
MDH 3
MDH 4
pGAN 5
pGAN 6
total
soluble
elution
kDA
180
130
100
70
55
40
35
25
15
10
pGAN 7
pGAN 8
pGAN 9
pGAN 10
pGAN 11
pGAN 12
pGAN 13
pGAN 14
pGAN 15
pGAN 17
pGAN 18
pGAN 19
pGAN 20
pGAN 21a
pGAN 22a

**D.**

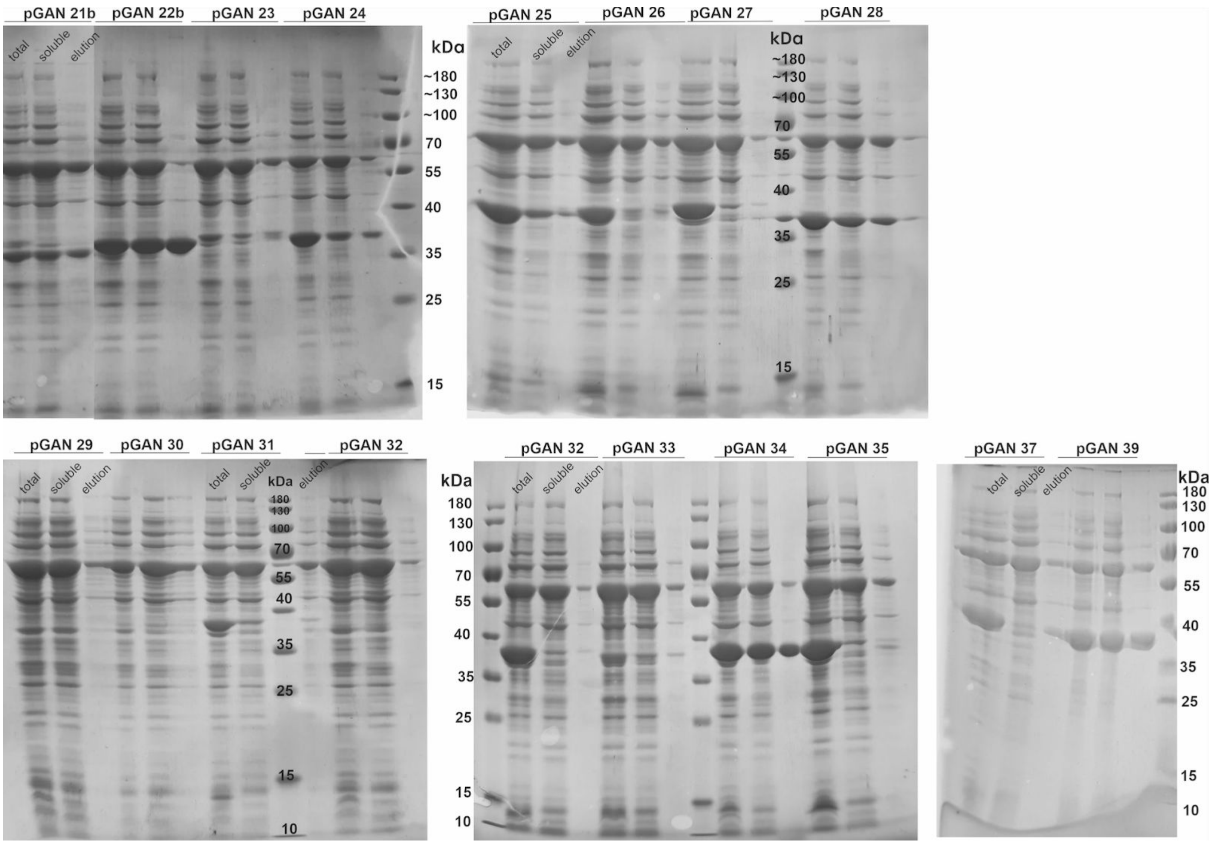
pGAN 21b
pGAN 22b
pGAN 23
pGAN 24
total
soluble
elution
kDa
~180
~130
~100
70
55
40
35
25
15
pGAN 25
pGAN 26
pGAN 27
pGAN 28
pGAN 29
pGAN 30
pGAN 31
pGAN 32
pGAN 33
pGAN 34
pGAN 35
pGAN 37
pGAN 39
180
130
100
10

**E.**

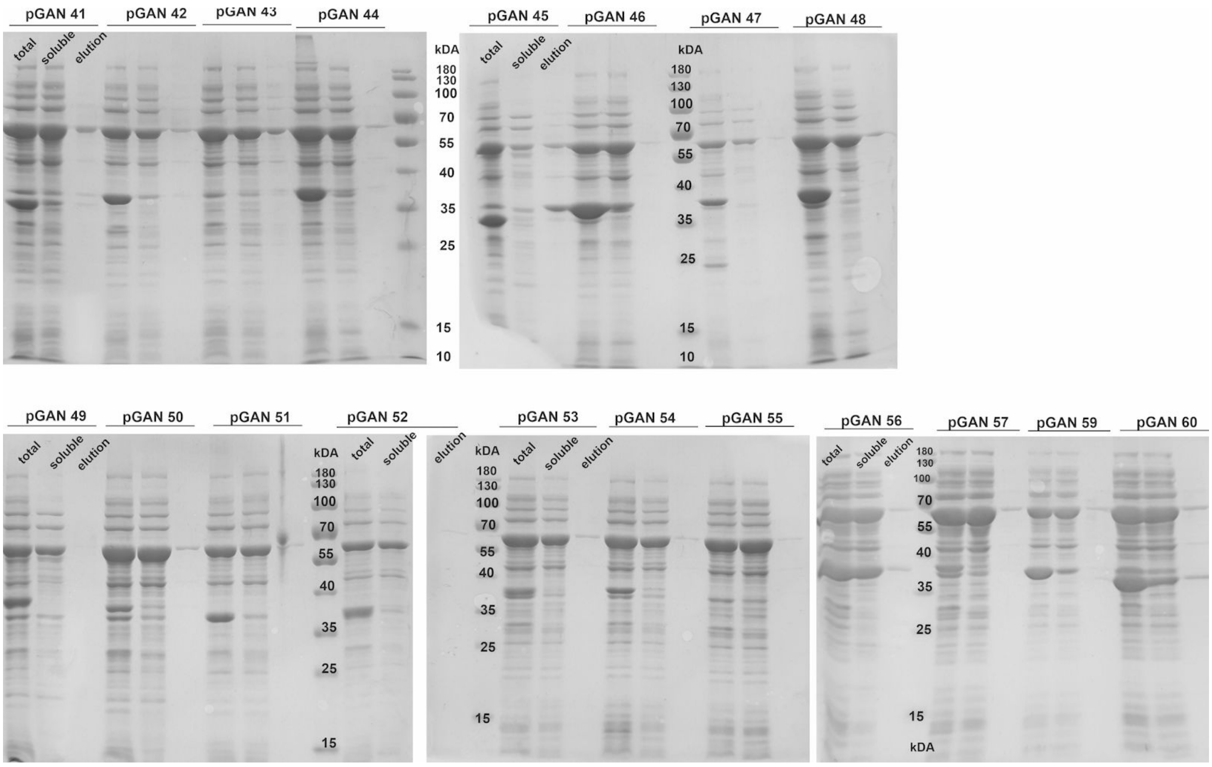

**Supplementary Figure 11 | SDS-PAGE gels of purified proteins.** (A) Batch 1, method 1 (see Methods). (B) Batches 2 and 3, method 1. (C) Batch 1, method 2. T – total lysate, S – soluble lysate, E – elution after affinity column use. (D) Batch 2, method 2. (E) Batch 3, method 2. The results are summarized in Supplementary Table 3.

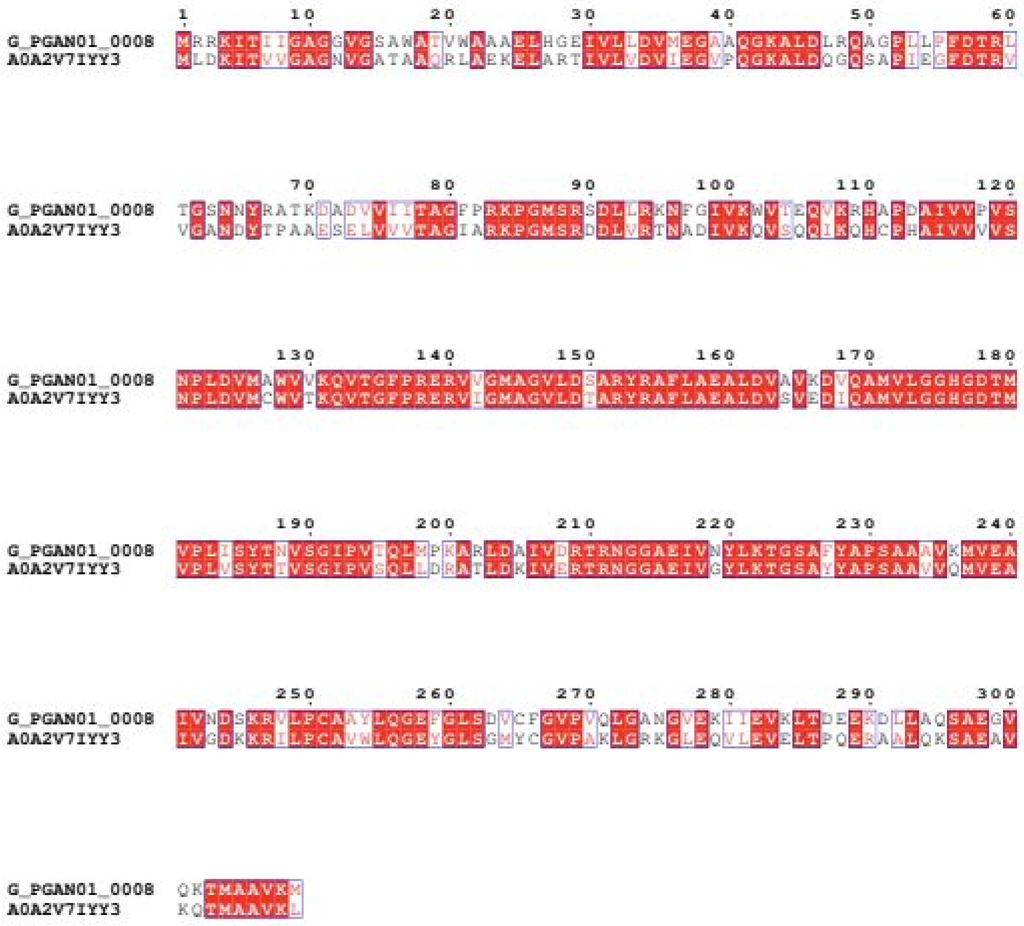

**Supplementary Figure 12 | Global sequence alignment of generated MDH enzyme (G_PGAN01_0008 aka pGAN 9) with its closest existing natural MDH enzyme from *Gemmatimonadetes bacterium* species (UniprotID: A0A2V7IYY3).** Generated enzyme contains 106 mutations comparing to its closest existing natural analog and display robust catalytic activity (Supplementary figure 14B, pGAN 9 identifier).

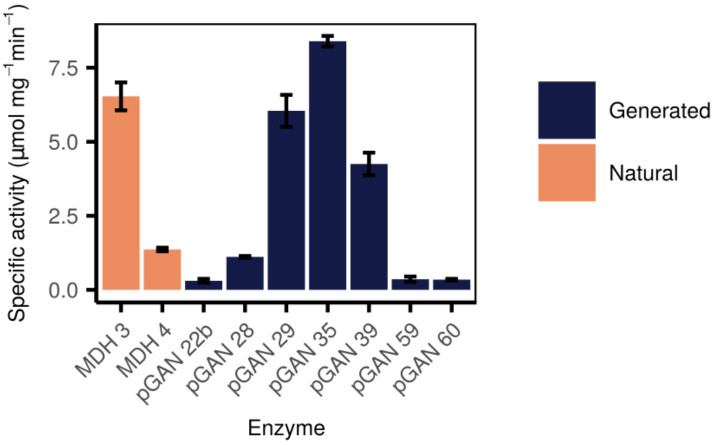

**Supplementary Figure 13 | Determined protein activities based on initial reaction rates as was measured in Method 1.**

**A.**

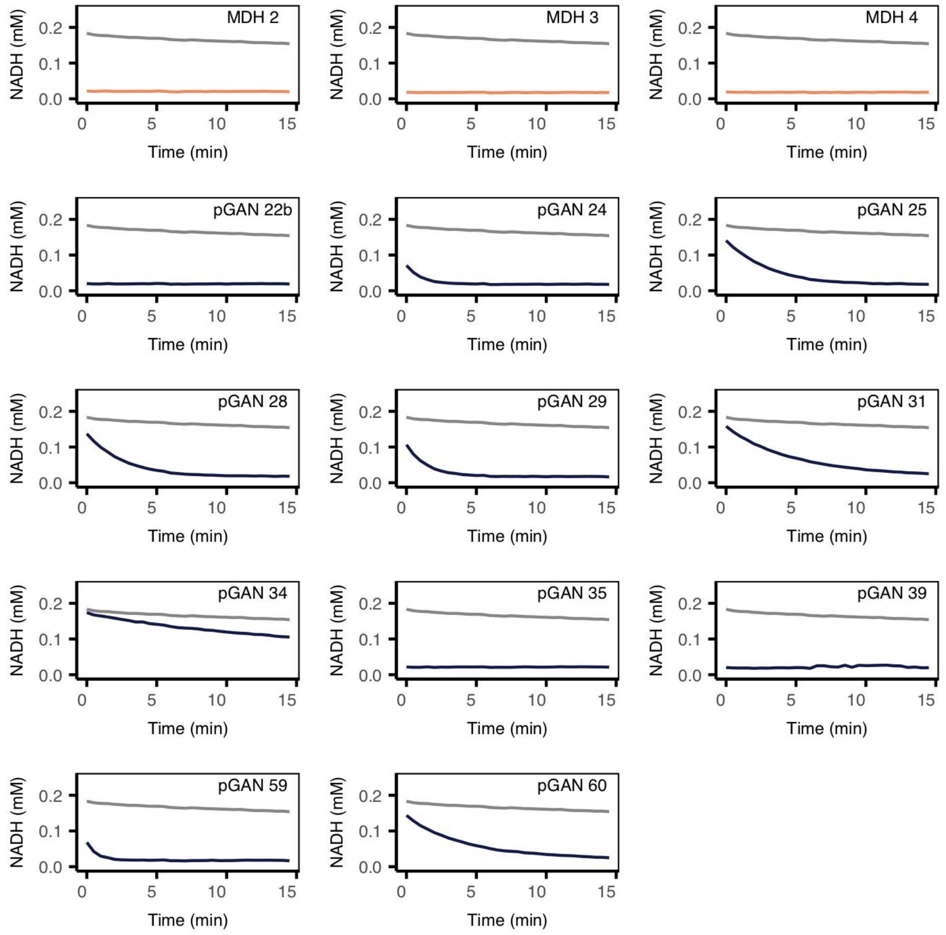

**B.**

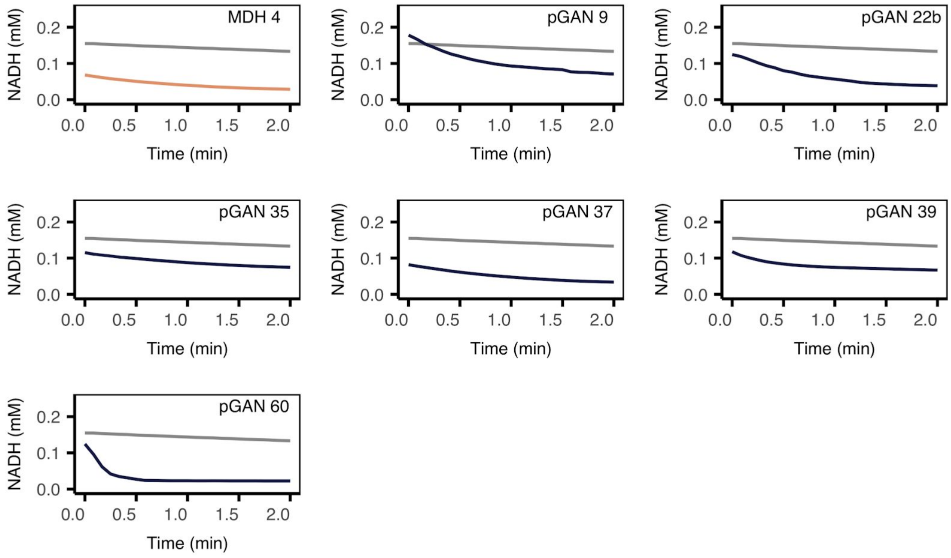

**Supplementary Figure 14 | Protein activities measured *in vitro* by monitoring NADH consumption.** (A) Activity data obtained for all soluble and active enzymes obtained using Method 1. An *E. coli* strain producing bovine serum albumin (BSA) was used as a negative control. Wild-type malate dehydrogenases (MDH 2, MDH 3, MDH 4), likewise produced in *E. coli.* serves as positive controls. (B) Activity data obtained for all soluble and active enzymes obtained using Method 2. An

*E. coli* strain producing green fluorescent protein (GFP) was used as a negative control. The wild-type malate dehydrogenase MDH 4 serves as a positive control. A time-dependent decrease in NADH can be readily observed for moderately active enzymes. For highly active enzymes all the NADH was consumed before measurements could take place, resulting in a horizontal line at the bottom of the plot. For clarity, each enzyme in the experiment is plotted in a separate panel using the same negative control data. Negative controls are plotted in grey, positive controls in orange, and generated sequences in blue.

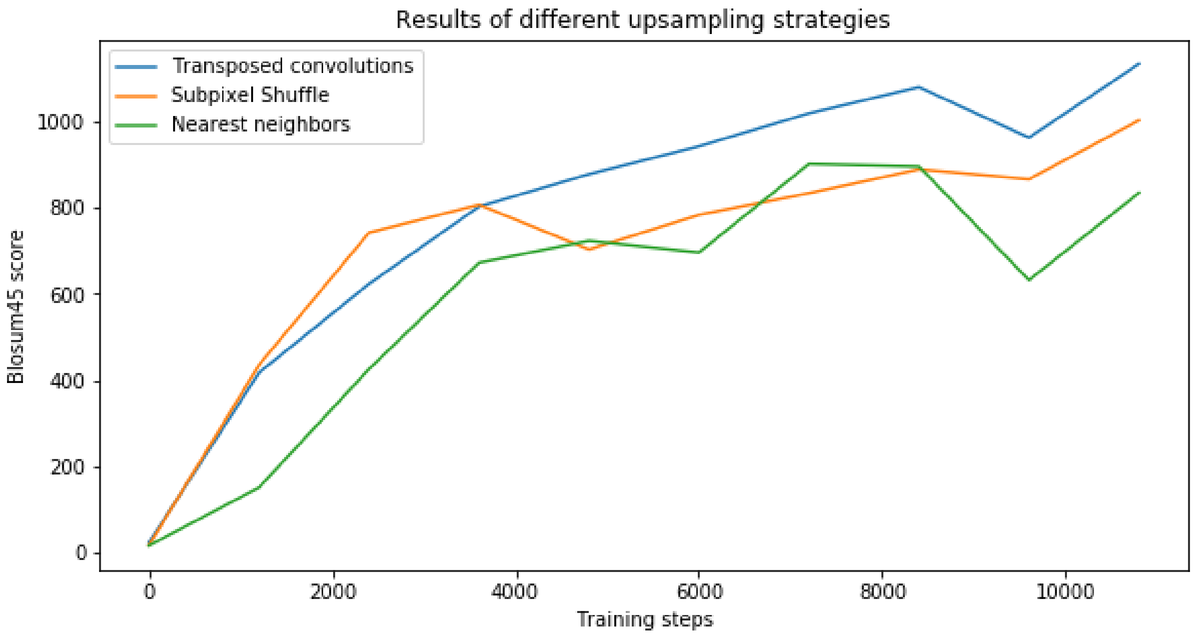

**Supplementary Figure 15 | GAN performance using different up-sampling strategies.** Model performances were measured using BLOSUM45 scores against training sequences for the first 10,800 steps. Blue line shows the BLOSUM45 scores of the final model that used transposed convolutions as the up-sampling technique. Orange line shows the results of the same model using sub-pixel shuffle, whereas green shows results of model that used nearest-neighbor interpolation to increase the dimensionality.

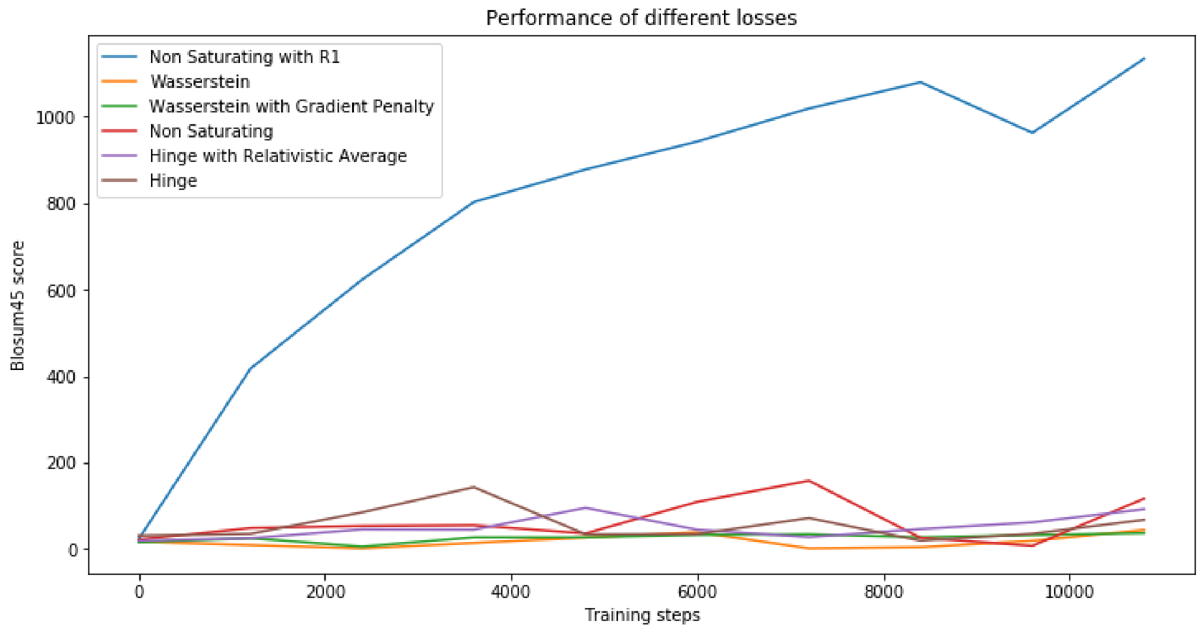

**Supplementary Figure 16 | GAN performance in first 10k steps using different losses.** Model performances were measured using BLOSUM45 scores against training sequences for the first 10,800 steps. Blue line shows BLOSUM45 scores of the final model that used non-saturating loss with R1 regularization. Orange line shows the results of the same model using the Wasserstein loss, whereas green shows results of the model that used the Wasserstein loss with a gradient penalty. We also experimented with original GAN loss [7] referred to as non-saturating (red line), hinge with relativistic average (purple line) and hinge loss (brown line).

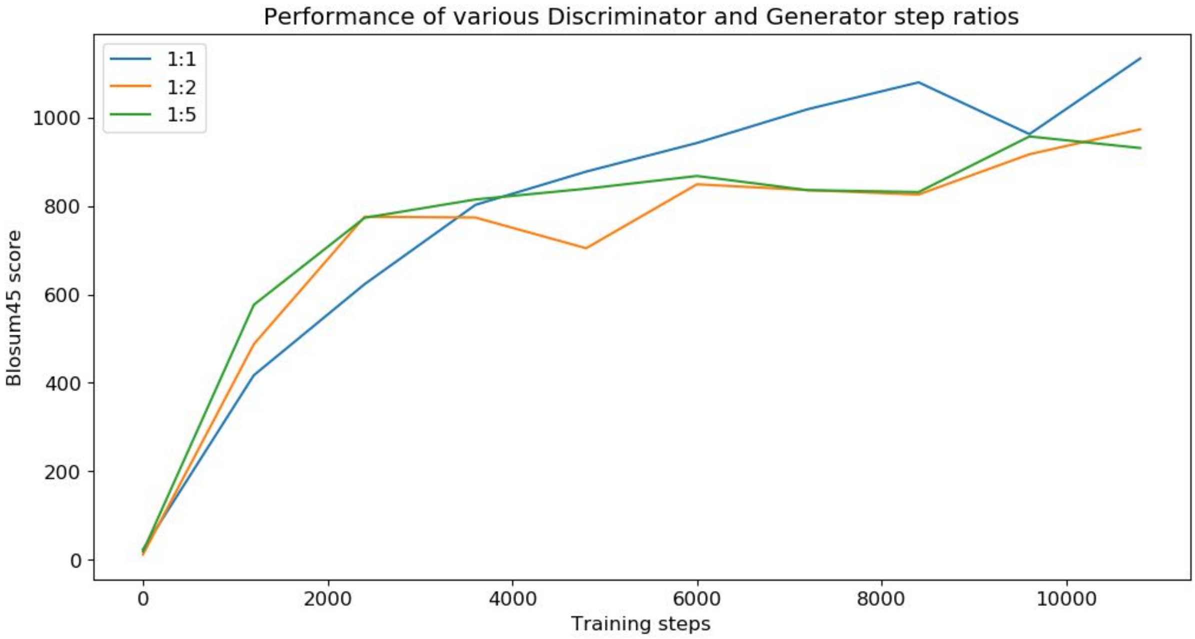

**Supplementary Figure 17 | GAN performance using different ratios between discriminator and generator steps.** Model performances were measured using BLOSUM45 scores against training sequences for the first 10,800 steps. Blue line shows the BLOSUM45 scores of the final model that used a 1:1 ratio. Orange and green lines show results of models that used 1:2 and 1:5 ratios, respectively.

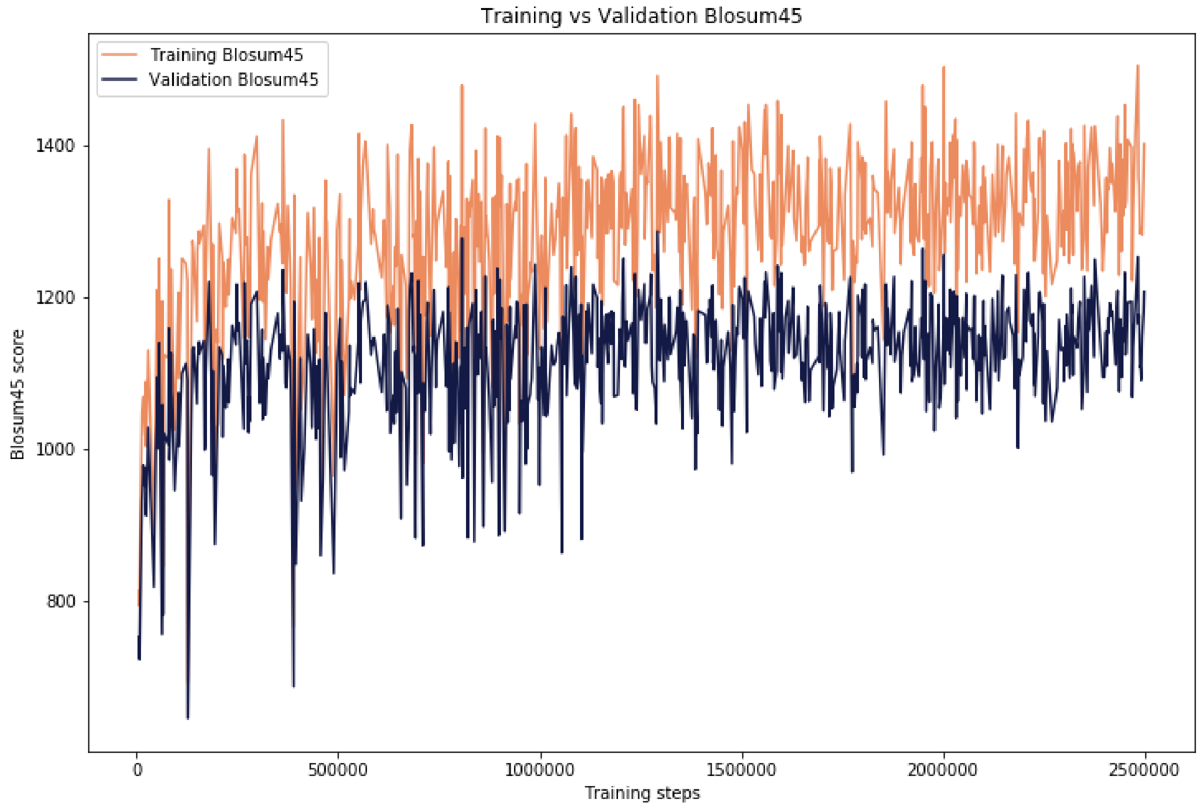

**Supplementary Figure 18 | Average BLOSUM45 scores of batches of generated sequences when compared with training and validation.** BLOSUM45 score was calculated using BLAST tool with BLOSUM45 matrix against sequences in the training and validation datasets separately.

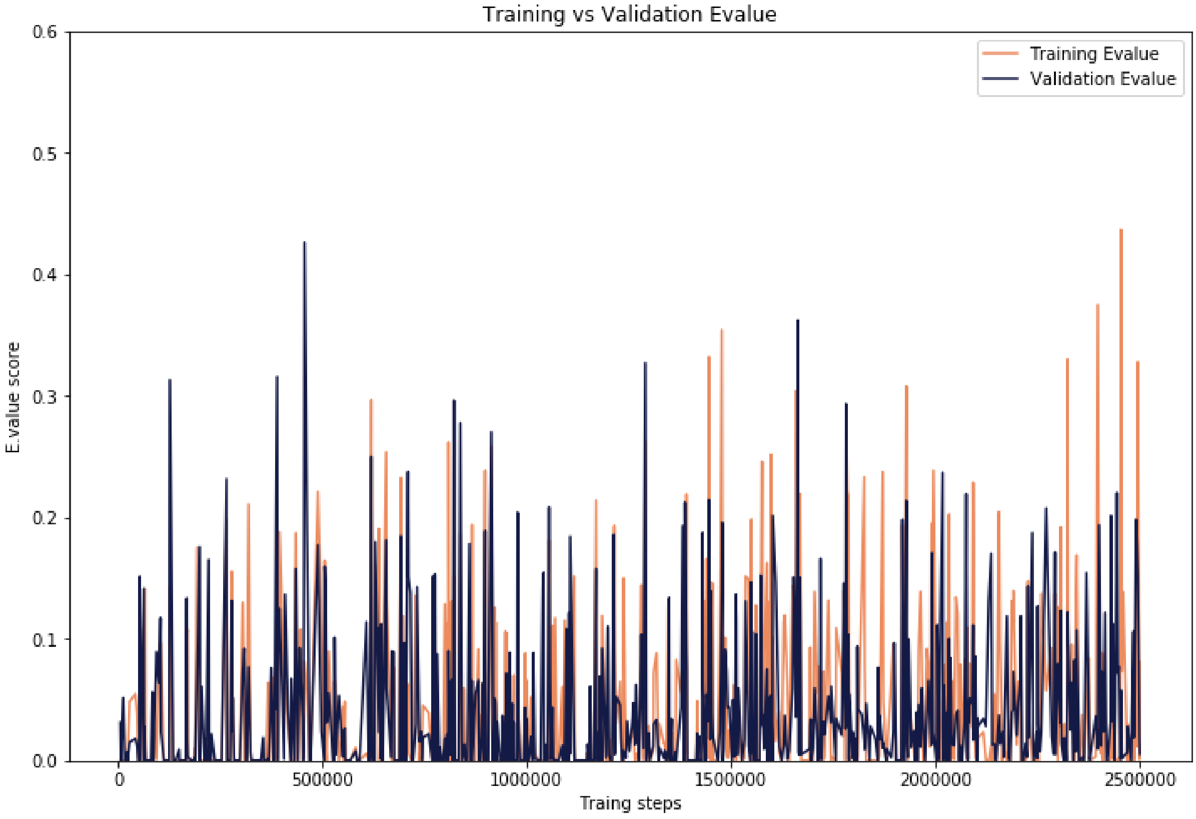

**Supplementary Figure 19 | Average *E*-values of batches of generated sequences when compared with training and validation sequences.** *E*-values were calculated using BLAST tool against sequences in the training and validation datasets separately.

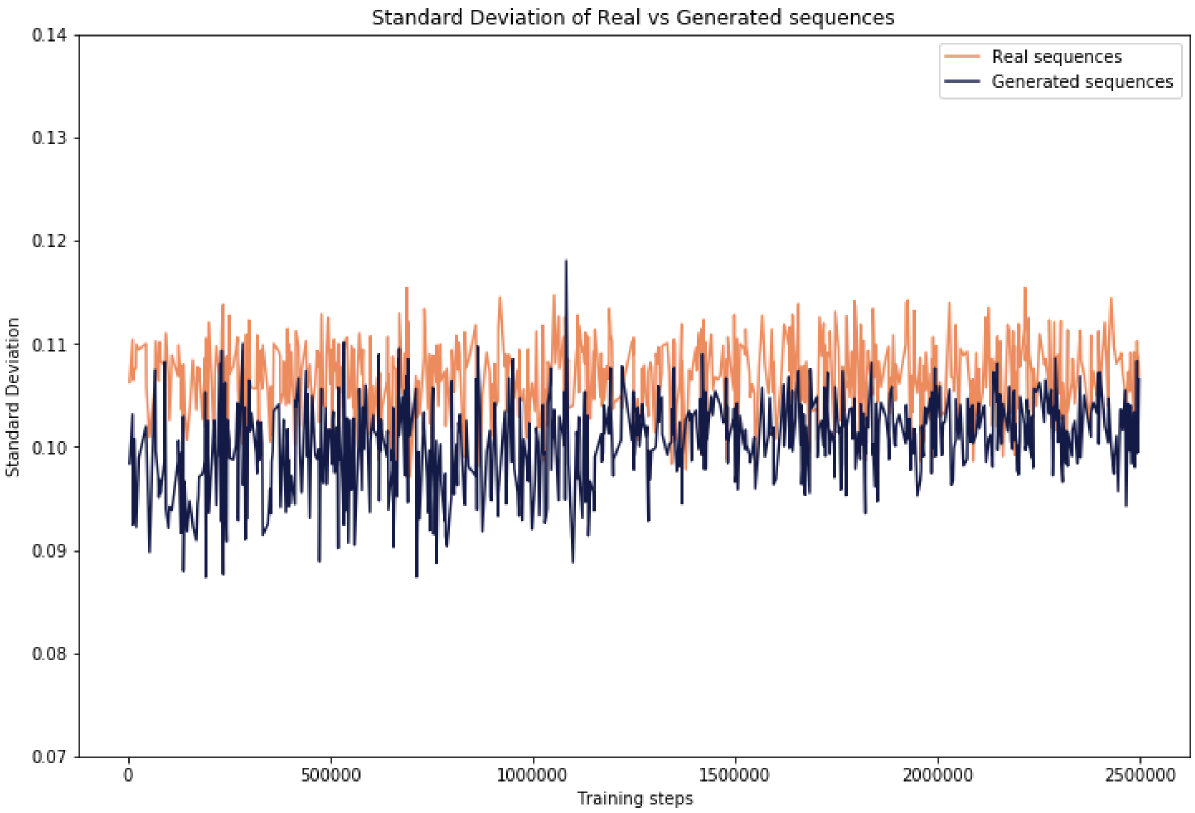

**Supplementary Figure 20 | Standard Deviation of the output of the last layer in the last Resnet block for real and generated sequences.** Low variation of generated sequences would indicate mode collapse. Having comparable variation of real and generated sequences indicates that the diversity of generated samples is similar to ones in the training dataset.

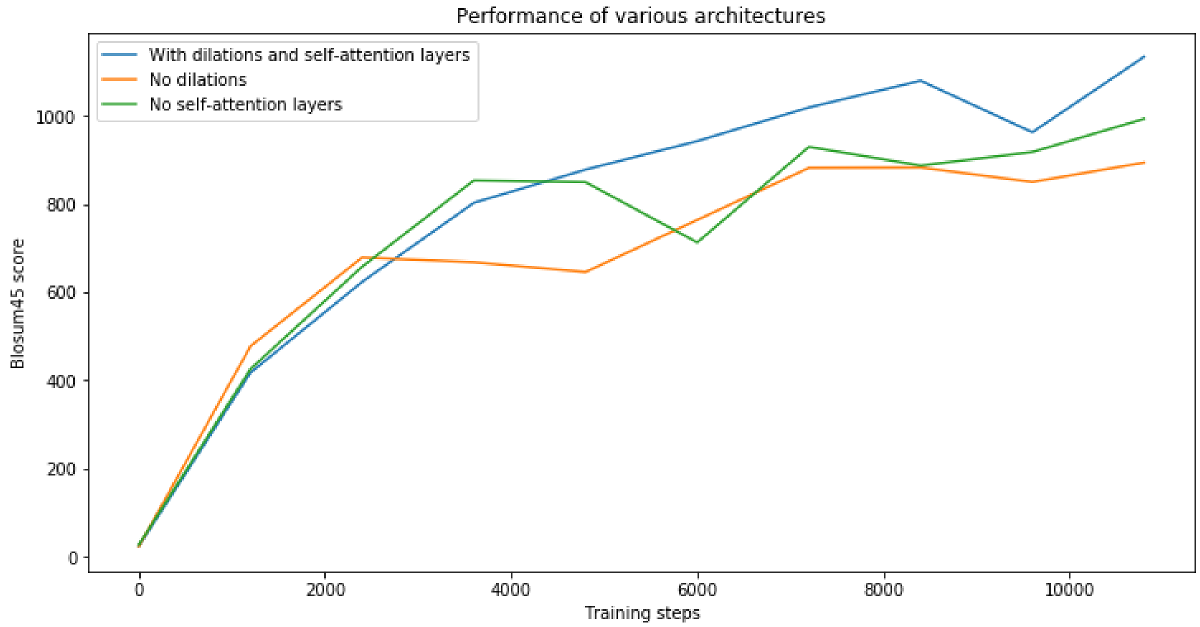

**Supplementary Figure 21 | Performance of self-attention and dilations layers.** Model performances were measured using BLOSUM45 scores against training sequences for the first 10,800 steps. Blue line shows the results of the final model that was used to produce sequences. It contained convolutional layers with dilations as well as self-attention layers. Green line shows the results of the same model without self-attention layers, whereas orange shows results of the model that did not use dilation in convolutional layers.

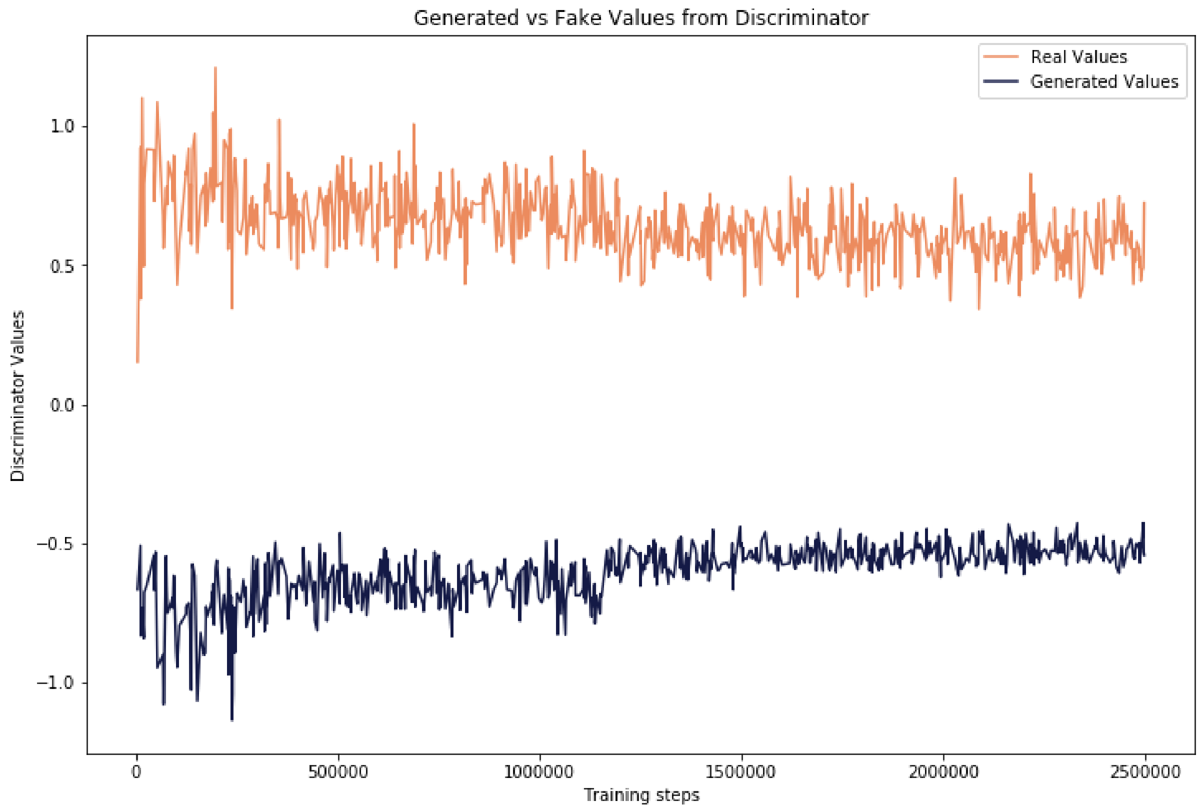

**Supplementary Figure 22 | Average values given by discriminator for real and generated sequences.** On average, the discriminator was able to distinguish between real and generated data well, which meant that the discriminator provided valuable feedback to the generator. This also indicates that the generator did not yet become good enough to fool the discriminator.

### Supplementary Tables

**Supplementary Table 1 | Similarities between distributions of individual amino acids and their groups with specific properties based on the Wilcoxon rank-sum test.** Only single amino acids have an identical distribution in both training and generated datasets (p>0.05), while multiple amino acid groups with specific physicochemical properties have identical distributions.

| Amino acids | Statistic | p-value | Properties |
|---|---|---|---|
| W | -44.5535 | 0 | - |
| T | -32.6757 | 3.45E-234 | - |
| N | -31.0134 | 3.55E-211 | - |
| P | 5.414836 | 6.13E-08 | - |
| F | 36.10193 | 2.12E-285 | - |
| A | -7.00421 | 2.48E-12 | - |
| G | 2.373175 | 0.017636 | - |
| I | 10.88373 | 1.38E-27 | - |
| L | 24.0913 | 3.08E-128 | - |
| H | 0.883687 | 0.376865 | - |
| R | 16.52561 | 2.40E-61 | - |
| M | 2.927137 | 0.003421 | - |
| V | -37.0289 | 3.93E-300 | - |
| E | -17.0094 | 7.00E-65 | - |
| Y | -2.82555 | 0.00472 | - |
| V, I, L, F, W, Y, M | -0.64435 | 0.519345 | Hydrophobic |
| S, T, H, N, Q, E, D, K, R | -3.965 | 7.34E-05 | Hydrophilic |
| F, W, Y, H | -1.00434 | 0.315217 | Aromatic |
| P, G, A, S | 7.263205 | 3.78E-13 | Small |
| K, R, H | 5.612656 | 1.99E-08 | Positive |
| D, E | -22.965 | 1.04E-116 | Negative |

| | | | |
|---|---|---|---|
| V, I, L, M | -2.04672 | 0.040685 | Aliphatic |
| S, C, T, M | -0.12037 | 0.904194 | Hydroxyl/sulfur |
| S, T, C, M, N, Q | -11.1188 | 1.02E-28 | Polar uncharged |
| H, K, R, E, D | 1.296764 | 0.194713 | Charged |

**Supplementary Table 2. Generated sequences and their identities to the closest real sequence.**

**Filename: 'Supplementary_table_identities.xlsx'**

**Supplementary Table 3 | Overview and assessment of experimental results.** Protein solubility was assessed based on SDS-page gels (Supplementary Figure 11) and activity was assessed based on measurements of MDH activity (Supplementary Figure 14).

| Method | Batch num. | Sample name | Solubility | Solubility comment | Activity | Activity comment | Final assessment |
|---|---|---|---|---|---|---|---|
| method 2 | batch 1 | MDH 2 | Pass | assessed purity above 80% | Fail | activity corresponds to GFP neg. control | |
| | | MDH 3 | Pass | assessed purity above 80% | Not tested | | |
| | | MDH 4 | Pass | assessed purity above 80% | Pass | wt pos. control | Pass |
| | | pGAN 5 | Fail | faint protein band - low purity | Not tested | | |
| | | pGAN 6 | Fail | no band corresponding to protein | Not tested | | |
| | | pGAN 7 | Fail | no band corresponding to protein | Not tested | | |
| | | pGAN 8 | Fail | no band corresponding to protein | Not tested | | |
| | | pGAN 9 | Pass | visible protein band - purity above 10%, possible protein clipping | Pass | | Pass |
| | | pGAN 10 | Fail | no band corresponding to protein | Not tested | | |
| | | pGAN 11 | Fail | no band corresponding to protein | Not tested | | |
| | | pGAN 12 | Fail | no band corresponding to protein | Not tested | | |
| | | pGAN 13 | Fail | faint protein band - low purity | Fail | activity corresponds to GFP neg. control | |
| | | pGAN 14 | Fail | no band corresponding to protein | Not tested | | |
| | | pGAN 15 | Pass | assessed purity above 50% | Fail | activity corresponds to GFP neg. control | |
| | | pGAN 17 | Fail | no band corresponding to protein | Not tested | | |
| | | pGAN 18 | Fail | no band corresponding to protein | Not tested | | |
| | | pGAN 19 | Fail | no band corresponding to protein | Not tested | | |
| | | pGAN 20 | Fail | no band corresponding to protein | Not tested | | |
| | | pGAN 21a | Fail | no band corresponding to protein | Not tested | | |

| | | | | | | | |
|---|---|---|---|---|---|---|---|
| | | pGAN 22a | Fail | no band corresponding to protein | Fail | activity corresponds to GFP neg. control | |
| | batch 2 | pGAN 21b | Pass | assessed purity above 30% | Fail | activity corresponds to GFP neg. control | |
| | | pGAN 22b | Pass | assessed purity above 80% | Pass | | Pass |
| | | pGAN 23 | Pass | assessed purity above 30% | Fail | activity corresponds to GFP neg. control | |
| | | pGAN 24 | Pass | assessed purity above 50% | Fail | activity corresponds to GFP neg. control | |
| | | pGAN 25 | Pass | assessed purity above 30% | Fail | activity corresponds to GFP neg. control | |
| | | pGAN 26 | Pass | assessed purity above 10% | Pass | higher absorbance than other proteins | |
| | | pGAN 27 | Fail | faint protein band - low purity | Fail | activity corresponds to GFP neg. control | |
| | | pGAN 28 | Pass | assessed purity above 30% | Pass | Too much NADH, pipetting error | |
| | | pGAN 29 | Fail | no band corresponding to protein | Not tested | | |
| | | pGAN 30 | Fail | no band corresponding to protein | Not tested | | |
| | | pGAN 31 | Fail | no band corresponding to protein | Pass | | |
| | | pGAN 32 | Fail | no band corresponding to protein | Not tested | | |
| | | pGAN 33 | Fail | no band corresponding to protein | Fail | activity corresponds to GFP neg. control | |
| | | pGAN 34 | Fail | faint protein band - low purity | Pass | | |
| | | pGAN 35 | Pass | assessed purity above 80% | Pass | | Pass |
| | | pGAN 36 | Fail | faint protein band - low purity | Not tested | | |
| | | pGAN 37 | Pass | assessed purity above 30% | Pass | | Pass |
| | | pGAN 39 | Pass | assessed purity above 50% | Pass | | Pass |
| | batch 3 | pGAN 41 | Fail | no band corresponding to protein | Fail | activity corresponds to GFP neg. control | |

| | | | | | | | |
|---|---|---|---|---|---|---|---|
| | | pGAN 42 | Fail | no band corresponding to protein | Not tested | | |
| | | pGAN 43 | Fail | no band corresponding to protein | Not tested | | |
| | | pGAN 44 | Fail | no band corresponding to protein | Not tested | | |
| | | pGAN 45 | Fail | band not corresponding to protein | Not tested | | |
| | | pGAN 46 | Fail | no band corresponding to protein | Not tested | | |
| | | pGAN 47 | Fail | no band corresponding to protein | Not tested | | |
| | | pGAN 48 | Fail | no band corresponding to protein | Not tested | | |
| | | pGAN 49 | Fail | no band corresponding to protein | Not tested | | |
| | | pGAN 50 | Fail | no band corresponding to protein | Not tested | | |
| | | pGAN 51 | Fail | no band corresponding to protein | Not tested | | |
| | | pGAN 52 | Fail | no band corresponding to protein | Not tested | | |
| | | pGAN 53 | Fail | no band corresponding to protein | Not tested | | |
| | | pGAN 54 | Fail | no band corresponding to protein | Not tested | | |
| | | pGAN 55 | Fail | no band corresponding to protein | Not tested | | |
| | | pGAN 56 | Pass | assessed purity above 50% | Fail | activity corresponds to GFP neg. control | |
| | | pGAN 57 | Fail | no band corresponding to protein | Not tested | | |
| | | pGAN 59 | Fail | no band corresponding to protein | Not tested | | |
| | | pGAN 60 | Pass | assessed purity above 30% | Pass | | Pass |
| method 1 | batch 1 | MDH 2 | Pass | assessed purity above 90%, dimers present | Pass | wt pos. control | Pass |
| | | MDH 3 | Pass | assessed purity at 100% | Pass | wt pos. control | Pass |
| | | MDH 4 | Pass | assessed purity above 90%, dimers present | Pass | wt pos. control | Pass |
| | | pGAN 5 | Fail | no band corresponding to protein | Not tested | | |

| | | | | | | | |
|---|---|---|---|---|---|---|---|
| | | pGAN 6 | Fail | no band corresponding to protein | Not tested | | |
| | | pGAN 7 | Fail | no band corresponding to protein | Not tested | | |
| | | pGAN 8 | Fail | no band corresponding to protein | Not tested | | |
| | | pGAN 9 | Fail | no band corresponding to protein | Not tested | | |
| | | pGAN 10 | Fail | no band corresponding to protein | Not tested | | |
| | | pGAN 11 | Fail | no band corresponding to protein | Not tested | | |
| | | pGAN 12 | Fail | no band corresponding to protein | Not tested | | |
| | | pGAN 13 | Pass | assessed purity at 100%, clipped protein | Fail | activity corresponds to BSA neg. control | |
| | | pGAN 14 | Fail | no band corresponding to protein | Not tested | | |
| | | pGAN 15 | Fail | no band corresponding to protein | Not tested | | |
| | | pGAN 17 | Fail | no band corresponding to protein | Not tested | | |
| | | pGAN 18 | Fail | no band corresponding to protein | Not tested | | |
| | | pGAN 19 | Fail | no band corresponding to protein | Not tested | | |
| | | pGAN 20 | Fail | no band corresponding to protein | Not tested | | |
| | | pGAN 21a | Fail | no band corresponding to protein | Not tested | | |
| | | pGAN 22a | Fail | no band corresponding to protein | Not tested | | |
| | batch 2 | pGAN 21b | Fail | no band corresponding to protein | Fail | activity corresponds to BSA neg. control | |
| | | pGAN 22b | Pass | assessed purity above 90%, dimers present | Pass | | Pass |
| | | pGAN 23 | Fail | no band corresponding to protein | Fail | activity corresponds to BSA neg. control | |
| | | pGAN 24 | Pass | assessed purity above 90%, protein clipping | Pass | | Pass |

| | | | | | | | |
|---|---|---|---|---|---|---|---|
| | | pGAN 25 | Pass | assessed purity above 90%, protein clipping | Pass | | Pass |
| | | pGAN 26 | Fail | no band corresponding to protein | Fail | activity corresponds to BSA neg. control | |
| | | pGAN 27 | Fail | no band corresponding to protein | Fail | activity corresponds to BSA neg. control | |
| | | pGAN 28 | Pass | assessed purity above 90%, protein clipping | Pass | | Pass |
| | | pGAN 29 | Pass | assessed purity above 90%, protein clipping | Pass | | Pass |
| | | pGAN 30 | Fail | no band corresponding to protein | Pass | | |
| | | pGAN 31 | Pass | assessed purity at 100%, faint band | Pass | | Pass |
| | | pGAN 32 | Fail | no band corresponding to protein | Fail | activity corresponds to BSA neg. control | |
| | | pGAN 33 | Fail | no band corresponding to protein | Fail | activity corresponds to BSA neg. control | |
| | | pGAN 34 | Pass | assessed purity at 100%, faint band | Pass | | Pass |
| | | pGAN 35 | Pass | assessed purity above 90%, dimers present | Pass | | Pass |
| | | pGAN 36 | Fail | no band corresponding to protein | Fail | activity corresponds to BSA neg. control | |
| | | pGAN 37 | Fail | no band corresponding to protein | Fail | activity corresponds to BSA neg. control | |
| | | pGAN 39 | Pass | assessed purity at 100% | Pass | | Pass |

| | | | | | | |
|---|---|---|---|---|---|---|
| | | pGAN 40 | Fail | no band corresponding to protein | Fail | activity corresponds to BSA neg. control |
| | batch 3 | pGAN 41 | Fail | no band corresponding to protein | Fail | activity corresponds to BSA neg. control |
| | | pGAN 42 | Fail | no band corresponding to protein | Fail | activity corresponds to BSA neg. control |
| | | pGAN 43 | Fail | no band corresponding to protein | Fail | activity corresponds to BSA neg. control |
| | | pGAN 44 | Fail | no band corresponding to protein | Fail | activity corresponds to BSA neg. control |
| | | pGAN 45 | Fail | no band corresponding to protein | Fail | activity corresponds to BSA neg. control |
| | | pGAN 46 | Fail | no band corresponding to protein | Fail | activity corresponds to BSA neg. control |
| | | pGAN 47 | Fail | no band corresponding to protein | Fail | activity corresponds to BSA neg. control |
| | | pGAN 48 | Fail | no band corresponding to protein | Fail | activity corresponds to BSA neg. control |
| | | pGAN 49 | Fail | no band corresponding to protein | Fail | activity corresponds to BSA neg. control |
| | | pGAN 50 | Fail | no band corresponding to protein | Fail | activity corresponds to BSA neg. control |
| | | pGAN 51 | Fail | no band corresponding to protein | Fail | activity corresponds to BSA neg. control |
| | | pGAN 52 | Fail | no band corresponding to protein | Fail | activity corresponds to BSA neg. control |
| | | pGAN 53 | Fail | no band corresponding | Fail | activity corresponds to |

| | | | | | | | |
|---|---|---|---|---|---|---|---|
| | | | | to protein | | BSA neg. control | |
| | | pGAN 54 | Fail | no band corresponding to protein | Fail | activity corresponds to BSA neg. control | |
| | | pGAN 55 | Fail | no band corresponding to protein | Fail | activity corresponds to BSA neg. control | |
| | | pGAN 56 | Pass | assessed purity above 90% | Fail | activity corresponds to BSA neg. control | |
| | | pGAN 57 | Fail | no band corresponding to protein | Fail | activity corresponds to BSA neg. control | |
| | | pGAN 59 | Pass | assessed purity above 90% | Pass | | Pass |
| | | pGAN 60 | Pass | assessed purity above 90% | Pass | | Pass |

**Supplementary Table 4 | Discriminator and Generator step ratio timings.** Training time was measured from the start of the training until the 10,800th step of the generator.

| Step ratio | Training time |
|---|---|
| 1:1 | 1h 22min |
| 1:2 | 2h 5min |
| 1:5 | 4h 14min |
